# ACtivE: Assembly and CRISPR-targeted *in vivo* Editing for Yeast Genome Engineering Using Minimum Reagents and Time

**DOI:** 10.1101/2022.07.15.500277

**Authors:** Koray Malcı, Nestor Jonguitud-Borrego, Hugo van der Straten, Urtė Puodžiūnaitė, Emily J. Johnston, Susan J. Rosser, Leonardo Rios-Solis

## Abstract

Among the numerous genetic tools developed for yeast, CRISPR/Cas system has been a widely used genome editing method thanks to its sophistication. However, CRISPR methods for yeast generally rely on pre-assembled DNAs and extra cloning steps to deliver gRNA, Cas protein, and donor DNA. These laborious steps might hinder its usefulness. Here, we propose a convenient, rapid, standardizable CRISPR method, named Assembly and CRISPR-targeted *in vivo* Editing (ACtivE), which only relies on *in vivo* assembly of linear DNA fragments for both plasmid and donor DNA construction. Thus, depending on the user’s need, these parts can be easily selected and combined from a repository, serving as a toolkit for rapid genome editing without any expensive reagent. The toolkit contains verified linear DNA fragments, which are easy to store, share and transport at room temperature, drastically reducing expensive shipping costs and assembly time. After optimizing this technique, eight ARS-close loci in the yeast genome were also characterized in terms of integration and gene expression efficiencies and the impacts of the disruptions of these regions on cell fitness. The flexibility and multiplexing capacity of the ACtivE were shown by constructing β-carotene pathway. In only a few days, > 80% integration efficiency for single gene integration and > 50% integration efficiency for triplex integration were achieved from scratch without using *in vitro* DNA assembly methods, restriction enzymes, or extra cloning steps. This study presents a standardizable method to be readily employed to accelerate yeast genome engineering and provides well-defined genomic location alternatives for yeast synthetic biology and metabolic engineering purposes.

## INTRODUCTION

Being a eukaryotic chassis, *Saccharomyces cerevisiae* has been extensively studied to produce high-value products from pharmaceuticals ^1–4^ to biofuels. ^5–8^ As a versatile and efficient genome engineering tool, the clustered regularly interspaced short palindromic repeats (CRISPR) system has been a widely used method to engineer yeast cell factories. ^9,10^

In the last decade, a great number of CRISPR-based methods have been developed for yeast genome editing or cell factory development. ^11,12^ Mainly, delivery or expression of CRISPR-associated (Cas) protein, guide RNA (gRNA), and donor DNA differ in these yeast-specific CRISPR methods. Generally, Cas protein (generally Cas9) which is responsible for the nuclease activity on a specific genomic location ^13,14^ is expressed through a plasmid vector ^15,16^ or genomic integration. ^17,18^ The latter needs an additional transformation for genomic integration and might lead to a burden on the host. gRNA forms a complex with Cas protein and guides it towards the target sequence ^13,14^ and it is generally expressed by using plasmid vectors in the yeast. A single plasmid containing the genes of Cas protein and gRNA ^16,19^ or separate independent plasmids for each can be used with an additional selective marker. ^20,21^ Donor DNA is used as a DNA repair template for homology-directed repair (HDR)^22^ after a double-strand break (DSB) is formed by Cas/gRNA complex. When it comes to delivery of donor DNA, single-strand oligos ^16^, double-strand oligos ^23^, or linear DNA fragments containing overlapping regions for *in vivo* assembly ^24^ can be used as well as linearized plasmids containing pre-assembled expression cassettes. ^25^

Even though high efficiencies in genome editing or pathway construction were achieved with these methods, further characterization is still needed to reach a full consensus. ^11,12^ On the other hand, the main obstacle in these methods is the requirement of *in vitro* DNA assembly and additional cloning steps to construct CRISPR plasmids. Golden Gate assembly ^26^, Gibson assembly^® 27^, or uracil-specific excision reaction-based cloning (USER cloning) ^28^ are the most widely used *in vitro* plasmid construction methods ^16,23,25,29–31^. For instance, Ronda et al. (2015) developed the CrEdit method standing for CRISPR/Cas9 mediated genome editing, using the USER cloning to construct the CRISPR plasmids. ^25^ In another study, a Csy4-assisted CRISPR method was used for multiple genome editing, and the plasmid constructs were cloned using Gibson Assembly. ^32^ However, the use of these techniques needs additional steps making the CRISPR process more labor-intensive and time-consuming. Moreover, these methods require relatively expensive DNA assembly kits ^33,34^ or specific type IIS restriction enzymes that might hinder design flexibility as the DNA fragment to be cloned might contain the recognition sites. Therefore, a collection of ready-to-use DNA parts skipping these pre-assembly steps can accelerate the genome editing process. Once this collection is obtained, it can also minimize the PCR deviations caused by the use of different DNA polymerases, which can be an important problem even though there are limited reports on this issue. ^35^ Finally, this toolkit based on verified DNA parts is more stable and practical to transport at room temperature for a prolonged period than glycerol stocks or agar slants. It can drastically reduce shipping costs while improving access to lower-income communities to contribute to synthetic biology’s democratization.

In addition to how CRISPR is carried out, its genome editing efficiency is another critical issue. The sequence features of CRISPR RNA (crRNA), which has a base complementarity to the target DNA ^36^ play a primary role in on-target efficiency and off-target specificity. ^37^ It has also been shown that gRNA expression might affect the CRISPR efficiency. ^38^ Apart from gRNA-dependent factors, target genomic loci might have an impact because of the chromatin structures. ^39^ Apel et al. (2017) found substantial variability in terms of integration efficiency and expression rate for 23 characterized genomic loci using green fluorescence protein (GFP) in *S. cerevisiae*. ^40^ Also, Wu et al. (2017) screened 1044 loci in the yeast genome, reporting important variations in RFP expression among the different loci tested. ^41^ Therefore, characterization of genomic regions in terms of CRISPR and gene expression efficiencies is crucial for identifying optimal target regions in the yeast genome. Especially for metabolic pathway construction involving many heterologous genes or extra copies of native genes ^2^, the use of efficient loci is quite significant to improve production yield.

To accelerate the yeast strain development process, more convenient CRISPR methods reducing time, labor, and cost are necessary, as well as identification and characterization of optimal genomic loci for chromosomal integration of constructs. In the present study, we developed a modular, convenient and standardizable CRISPR method, named named Assembly and CRISPR-targeted *in vivo* Editing (ACtivE), relying on *in vivo* assembly of co-transformed DNA modules in yeast. We used chemically synthesized gRNA cassettes and synthetic overlapping fragments as connectors for *in vivo* DNA assembly. In this way, the modules can be easily selected and combined from a part repository depending on the application. Particular parts such as a module expressing a specific type of Cas protein or a gRNA module targeting a specific genomic region can be combined along with other modules to *in vivo* construct the CRISPR plasmid. As the part repository contains only PCR-verified linear fragments, the use of plasmid purification kits to obtain the parts or extra enzymatic treatment steps such as DpnI digestion to degrade parental plasmid is not necessary. Without the use of agar stab/plate, this approach can also facilitate shipping. Also, compatible, and custom-made parts can be exchanged between different groups thanks to the connectors (synthetic fragments) at the terminals of each module. This collection can serve as a CRISPR toolkit that will be expanded with new modules and is freely available at https://www.leorioslab.org/cost-crispr-toolkit/ so that users will be able to perform yeast genome editing by simply providing their own custom donor DNA. Therefore, ACtivE allows rapid and plasmid-free genome engineering in the yeast genome as it abolishes *in vitro* DNA assembly and cloning steps so that it does not require the use of type II restriction enzymes or DNA assembly kits which can be a considerable expense for lower-income laboratory settings. After optimizing the genome editing efficiency and reproducibility of the method, eight different genomic regions were characterized using a GFP, mNeonGreen, in terms of determining integration and gene expression efficiencies as well as integration effects on cell fitness. ∼80% single gene integration and deletion efficiencies were achieved. Furthermore, the multiplexing capacity of ACtivE was tested for simultaneous integration of multi genes into multi loci in the genome. The CRISPR technique used in this study enables standardizable and rapid genome engineering in the yeast genome, and thoroughly characterized genomic regions provide more alternatives to be used for genomic integration or pathway construction.

## MATERIALS & METHODS

### Oligonucleotides, Reagents, and Plasmids

All primers used in the study are listed in Table S1-S4. The primers were ordered from Integrated DNA Technologies (IDT) as standard DNA oligos for fragments from 20 bp to 100 bp or as DNA ultramers for fragments with ∼120 bp length. Synthetic gRNA cassettes (Figure S1) were ordered from Twist Bioscience. Phusion Flash High-Fidelity PCR Master Mix (Thermo Fisher Scientific) and PrimeSTAR® GXL DNA Polymerase (TaKaRa) were used for PCR reactions while DreamTaq Green PCR Master Mix (Thermo Fisher Scientific) was used for colony PCR. FastDigest DpnI (Thermo Fisher Scientific) was used to degrade the parental plasmids. GeneJET PCR Purification Kit (Thermo Fisher Scientific) was used for PCR clean-up. p426_Cas9_gRNA-ARS511b (Addgene) was used to amplify uracil auxotrophic selection marker (*URA3*), yeast origin of replication (2μ ori) and bacteria storage fragment containing ampicillin resistance gene (*AmpR*), and bacterial origin of replication. pWS158 (Addgene) was used as a template to amplify *Cas9* (*Streptococcus pyogenes*) codon-optimized for expression in *Saccharomyces cerevisiae*. mNeonGreen was used as green fluorescence protein (GFP) and it was amplified from pCPS1ULA-BA6 was obtained as a gift from Matthew Dale (Rosser Lab, the University of Edinburgh). pTDH3-Re2.8-2 was gifted by Jamie Auxillos (Chris French Lab, the University of Edinburgh) and red fluorescent protein (RFP), mCherry, was amplified using this plasmid.

### Strains and Media

The parent strain of *S. cerevisiae*, BY4741 {MATa; *his3*Δ1; *leu2*Δ0; *met15*Δ0; *ura3*Δ0}, was used for genomic integrations and was kindly provided by Dariusz Abramczyk (Chris French Lab, the University of Edinburgh). *S. cerevisiae* CEN.PK2-1C {MATa; *his3*Δ1; *leu2*-3_112; *ura3*-52; *trp1*-289; *MAL2*-8c; *SUC2*} from EUROSCARF Collection was used for genomic deletions. Unless otherwise stated, all chemicals were sourced from Sigma-Aldrich at the highest available purity. For cultivation of strains, YPD medium containing yeast extract (1%(w/v)), peptone (2%(w/v)), and 2% (w/v) dextrose (glucose) was used. To select positive transformants expressing *URA3* marker, synthetic defined medium containing complete supplement mixture minus uracil (CSM-Ura, MP Biomedicals™), 0.17% (w/v) yeast nitrogen base without amino acid, 0.5% (w/v) ammonium sulphate, 2% (w/v) glucose, and 2% (w/v) agar was used. For counter-selection of plasmid-free yeast cells, a synthetic defined medium supplemented with 0.1% (w/v) 5-Fluoroorotic Acid (5-FOA) (Thermo Fisher Scientific) was used. YPD media and complete synthetic defined (SD) media containing all amino acids were used for mNeonGreen expression and characterization of genomic loci.

### Yeast Heat-Shock Transformation

The chemicals were sourced from Sigma-Aldrich unless otherwise stated. All transformations were carried out according to LiAc/PEG heat-shock method ^42^ with some small modifications. After overnight cultures, fresh cultures were prepared to obtain the cells in the exponential phase. The cells were then washed once and were pelleted by centrifugation. The transformation mix containing 240 μL PEG (50%(w/v)), 36 μL 1.0 M lithium acetate (LiAc) and 50 μL single-stranded carrier DNA (2.0 mg/mL) (herring sperm DNA, Promega) were added onto the cell pellet. Next, DNA fragments and water were added until the volume was made up to 360 μL. 50 fmol equivalent molarity of each plasmid-forming DNA part, 500 ng – 1000 ng from each donor DNA part were added to the transformation mixes. As a large number of modular fragments were used for multiplexing, transformation volume was increased to 400 μL when needed by adding more DNAs without water addition. After homogenous transformation mixes were obtained, the cells were incubated for 45 minutes at 42 °C. After plating cells to the selective media, the cells were incubated for 2-3 days at 30 °C.

### Determination of Genome Editing Efficiency

When the colonies became visible after integrating mNeonGreen into single loci, the plates were imaged on a blue-LED transilluminator (Thermo Fisher Scientific) to distinguish fluorescent mNeonGreen-expressing positive colonies from non-fluorescing negative colonies. Also, colony PCR was performed on randomly selected five positive colonies from each plate to confirm that the genes are integrated into correct locations, and 100% consistency was observed for all positive colonies controlled. Integration efficiency was determined by calculating the percentage of green light-emitting colonies. To count the colonies on the plates when numerous colonies were obtained, ImageJ ^43^, a free distribution software, was employed with the Colony Counter plugin (Figure S4). ^44^ The plate images on the blue-LED transilluminator were first converted to 16-bit pictures. The green colonies (positive) and white colonies (negative) were distinguished using color contrast, and colonies were counted automatically. For integration efficiency of simultaneous integration of mNeonGreen and mCherry, first, the mNeonGreen expressing colonies were determined on a blue-LED transilluminator. Those colonies were then screened in CLARIOstar Plus microplate reader (BMG Labtech) to detect mCherry expression using spectral scanning with an emission wavelength ranging from 580 nm to 670 nm at 552 nm excitation wavelength. The positive colonies expressing both mNeonGreen and mCherry were also screened by colony PCR to confirm the integrations. Finally, orange colonies were counted using Colony Counter – ImageJ to determine the integration efficiency of β-carotene pathway. These integrations were also confirmed by employing colony PCR. The genomic deletions were first screened using colony PCR, and the deletions were confirmed by Sanger sequencing performed at GENEWIZ, Inc (Leipzig, Germany).

### Characterization of Genomic Loci with mNeonGreen Expression

Three individual colonies from each strain expressing mNeonGreen on different loci were selected after confirming the integrations. 5-FOA counter-selection was performed to select plasmid-free cells after overnight culture in YPD media. Eight mNeonGreen expressing strains and the parent strains, BY4741, were then cultured in YPD and SD media with three replicates for 72 hours to observe expression rates of mNeonGreen on each locus. Biomass of different strains was also measured to compare the integration effect on cell fitness. After overnight culture of each strain, cells were inoculated into fresh media to be grown for around six hours to obtain cells in the exponential phase. The initial OD600 was adjusted to 0.1 for the growth experiments for all strains. To avoid sedimentation in low volumes, the cells were grown in 1 mL media, YPD or SD, using 24-well plates (Greiner) with shaking at 200 rpm, at 30 °C. To measure fluorescence intensity and biomass, 20 μL culture samples were taken on the 0^th^, 3^rd^, 6^th^, 12^th^, 15^th^, 18^th^, 21^st^, 24^th^, 36^th^, 48^th^, and 72^nd^ hours and mixed with water in 200 μL total volume in a black, clear-bottom 96-well plate (Greiner). An anti-condensation solution containing 0.05% Triton X-100 (Sigma-Aldrich) in 20% ethanol ^45^ was used to cover the lids of the well-plates to prevent condensation on the lids. Fluorescence intensities and OD_600_ measurements were taken using the CLARIOstar microplate reader (BMG Labtech). Matrix scan (2×2, 25 flashes) was used to scan the wells. To measure mNeonGreen expression, 490 nm and 525 nm wavelengths were used for excitation and emission, respectively, with 10 nm bandwidth. The emission wavelength was set to 585 nm for autofluorescence of media and yeast cells. ^46^ The gain was 1500 for both protocols.

### Data Analysis and Software

CRISPR experiments were conducted in at least three replicates. The error bars represent the standard deviations of different experiments. The one-way analysis of variance (ANOVA) (*p-*value<0.05) was used to determine whether there was a statistically significant difference between the experiments. The yeast genome was screened using UCSC Genome Browser ^47^ (http://genome.ucsc.edu) to find ARS-close intergenic regions. The potential crRNA sequences on each region were scored using CRISPOR ^48^ (http://crispor.tefor.net), an online gRNA selection tool giving sequence-based scores using sequence prediction algorithms. ^49,50^ The linear regression was used to determine the relationship between integration efficiencies and gRNA efficiency prediction scores, and Person’s correlation coefficients (R) were calculated employing MATLAB. The fluorescence intensity and OD600 data were analyzed using SciPy package ^51^ (Matplotlib, NumPy, pandas), *seaborn*, ^52^ and *omniplate* ^53^ in Python. Before analyzing growth characteristics, OD correction was performed using a standard curve for 2% (w/v) glucose-containing media as there is a non-linear relationship between biomass and OD_600_. To find the actual fluorescence intensity of per mNeonGreen expressing cell, autofluorescence caused by yeast cells themselves and media, YPD or SD, was corrected. ^46^ To calculate the areas under the curves, the trapezoidal rule was used. The codes and the standard curve used for plate reader data analysis can be found on https://github.com/kmalci/plate-reader. The illustrations were made using the BioRender. ^54^

## RESULTS & DISCUSSION

### Standardizable, Rapid, Convenient Yeast CRISPR

To create a CRISPR plasmid expressing the gRNA and Cas protein, *in vitro* DNA assembly procedures typically followed by a cloning step into *Escherichia coli* are the standard methods used in the yeast genome engineering processes. However, these steps retard this process, especially if sequential genome editing studies are needed to construct or design synthetic metabolic pathways. Here, we developed a more convenient, modular, and rapid CRISPR method, named ACtivE, which relies only on amplifying the functional units (donor DNA) through PCR. The modules contain small overlapping sequences assembled via *in vivo* HDR, so time-consuming and labor-demanding extra steps are omitted.

Gibson et al. (2008) succeeded in assembling a long, ∼600 kb, synthetic genome consisting of 25 overlapping fragments in yeast through HDR. ^55^ After this, Kuijpers et al. (2013) reported that using synthetic overlapping sequences increases the efficiency of *in vivo* DNA assembly in yeast. ^56^ In our design, we used five modules, a *Cas9* expression cassette, a gRNA expression cassette, a selection marker, a storage part, and a yeast origin of replication (ORI), to create an all-in-one CRISPR plasmid. 60 bp synthetic fragments ^56^ were used as overlapping sequences (connectors) between each adjacent module. Among them, selection marker and ORI are essential parts for surviving and propagating the plasmid, while Cas protein and gRNA are responsible for CRISPR activity. The storage part, an optional module, contains a bacteria-specific selection marker and ORI to be used for storage of the assembled plasmid in *E. coli* if desired. As chemical DNA synthesis is a rapidly growing sector ^57^ with many alternative approaches ^58^, synthetic production of expression cassettes is a reasonable and affordable choice, especially for short fragments, 300 bp – 500 bp, compared to traditional constructing methods. Therefore, synthetic gRNA expression cassettes were used in our method, allowing the users to choose any integration locus they desire in addition to the eight ones characterized in this work. When it comes to donor DNA to integrate heterologous genes of interest (GoI), we used four parts, ∼1000 bp upstream homology arm (UHA), promoter, CDS + terminator, and ∼1000 bp downstream homology arm (DHA). In this way, individual promoter parts could be easily changed as a widely used approach for fine-tuning gene expression. ^59^ Similar to plasmid assembly, each donor part had a 60 bp overlapping sequence with its adjacent part. Figure 1 illustrates the working principles and design of ACtivE and the content of the toolkit provided. Once plasmid modules are produced, they can be stored for subsequent study. Indeed, one of the benefits of this design is the CRISPR plasmid construction can be readily standardized by using the connectors (synthetic fragments) and a part collection containing different alternatives for each module. For instance, the Cas cassette has connector A and connector E at the terminals. Depending on the purpose, a particular type of Cas protein, Cas9, dead Cas9, Cas12a, etc., could be selected from the collection to be combined with other modules. This also applies to other parts, such as the selection markers, allowing more flexibility and part exchange between different research groups. Moreover, as the method relies on only *in vivo* assembly and PCR for the donor DNA, the whole process can be finished in a single day with a good organization from scratch.

**Figure 1:**
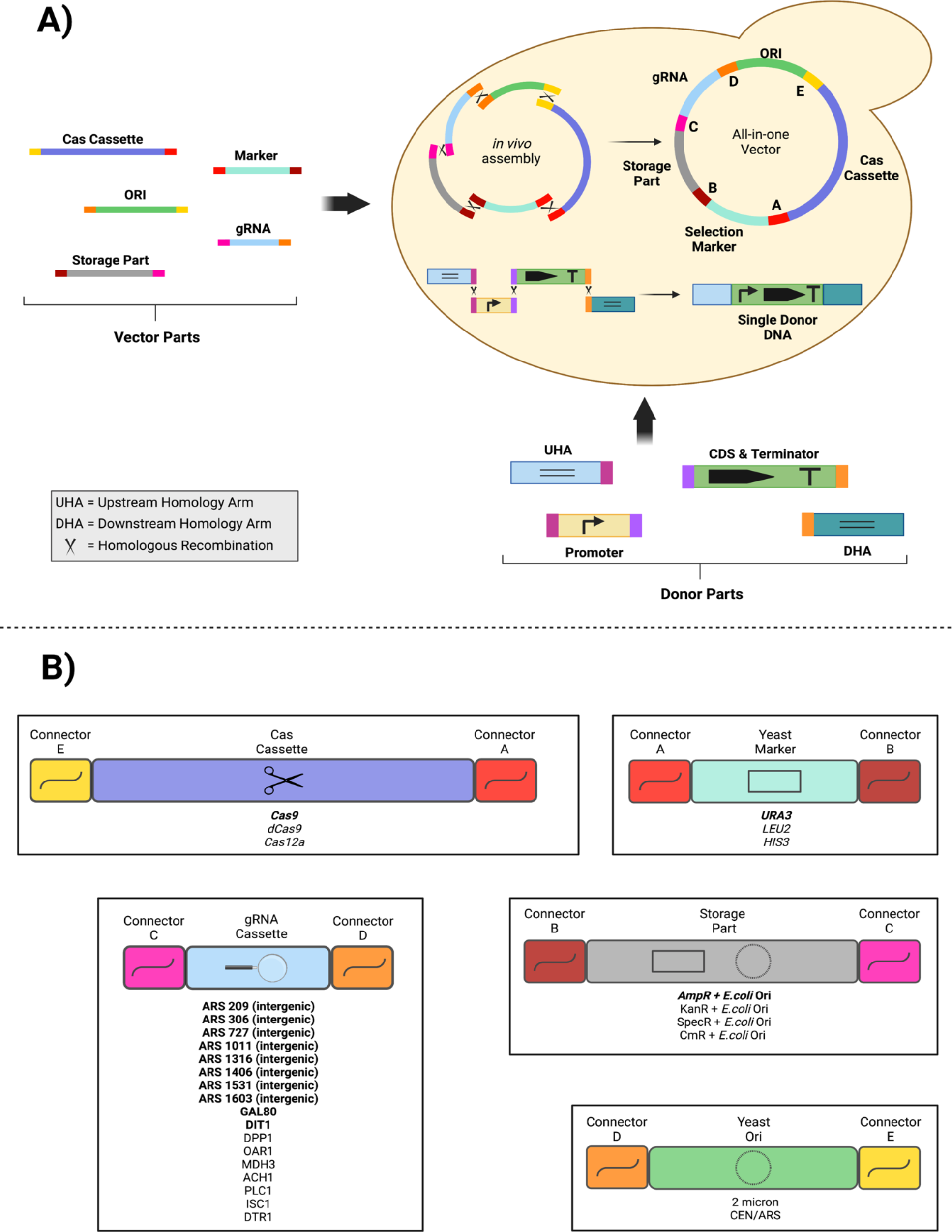
The overview of the ACtivE method. **A)** Each plasmid module is produced by PCR and can be stored to create a part collection. The modules contain an overlapping sequence (connector) with their neighbor fragment for *in vivo* assembly via homologous recombination. Depending on the application, the plasmid modules can be selected from the part collection. In this study, the donor DNA consisted of four parts, and they were co-transformed with plasmid modules in a single transformation step. In yeast, the parts are assembled via homologous recombination, and the donor DNA is inserted into the genome thanks to its homology arms with the genome. The length of the parts may not represent the actual sizes. **B)**The parts provided in the first version of the ACtivE toolkit (the parts in bold were characterized in this study). The gRNAs of ARS-close regions target intergenic regions in the genome whereas other gRNAs target the genes. By providing their own custom repair donor DNA, the users can readily combine the plasmid modules depending on their applications for a rapid yeast genome editing study. The toolkit will be expanded with new parts and is freely available with more information on https://www.leorioslab.org/cost-crispr-toolkit/

### Selecting Integration Regions

To integrate the gene of interest (GoI), *mNeonGreen*, into the yeast genome, eight loci on eight different chromosomes were selected to compare the integration and gene expression efficiency and impact on cell fitness. Previous studies reported that gene expression rates of heterologous genes tend to be higher if they are located in a region close to autonomously replicating sequences (ARS). ^60,61^ Therefore, ARS-close intergenic regions were targeted for integrations, and crRNA sequences on these regions were scored using different algorithms. Moreno-Mateos et al. (2015) developed a gRNA activity prediction algorithm, CRISPRscan, using zebrafish-specific gRNAs, giving higher efficiency scores if the crRNA sequence has high guanine but low adenine content. ^50^ Whereas Doench et al. (2016) used mammalian cells to develop a gRNA efficiency algorithm. ^49^ Both algorithms were considered for selecting crRNAs as yeast-specific algorithms have not been developed yet. To minimize the off-target effects in the genome, the sequences that have the maximum MIT scores representing the highest uniqueness were selected. ^62^

Table 1 shows the crRNA sequences used in this study and their corresponding scores. The sequences were selected considering the highest specificity and efficiency prediction scores among the other crRNA sequences on the regions.

**Table 1:**
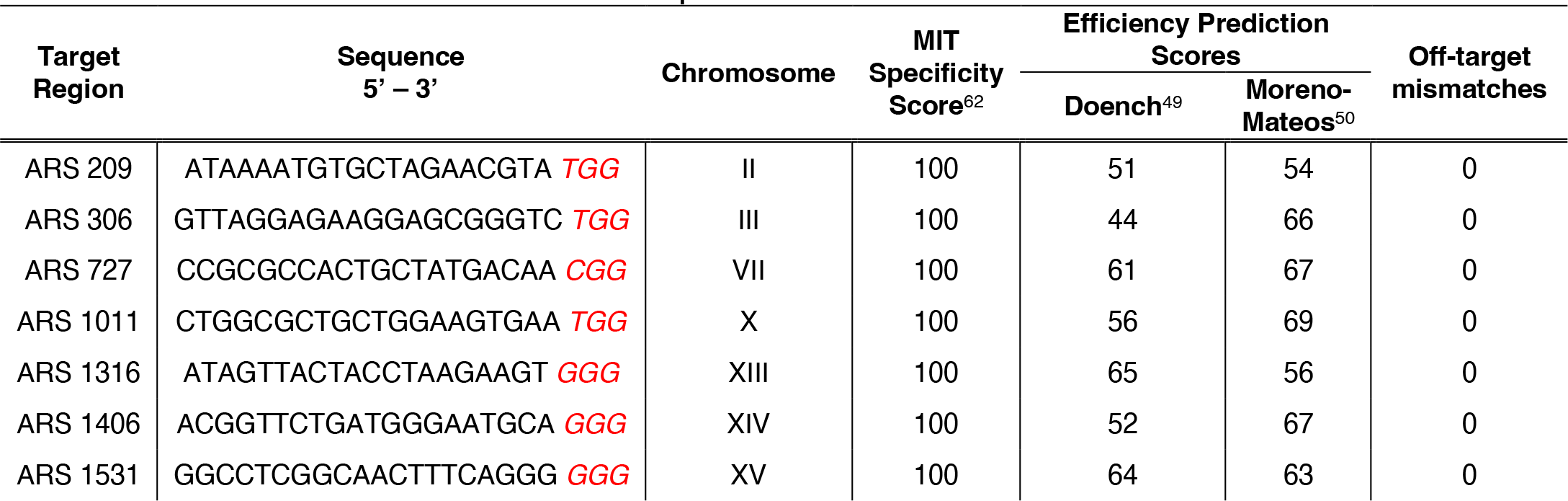
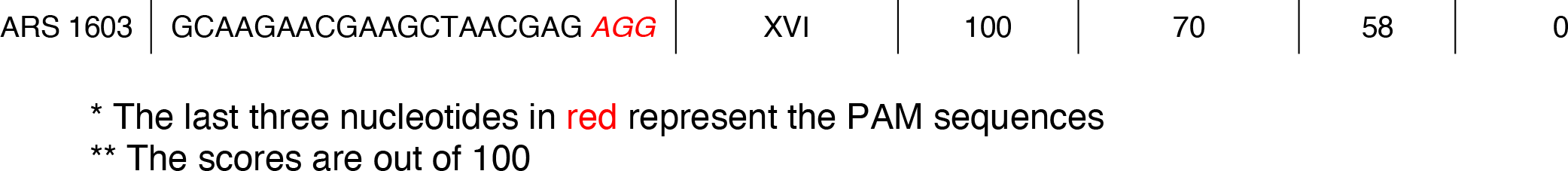
The crRNA sequences used in this study to integrate *mNeonGreen* and sequence features

### Optimization of the Method

The integration efficiency of ACtivE was first tested on three loci, ARS 306, ARS 1316, and ARS 1603, using high-fidelity Phusion polymerase to amplify plasmid modules and the donor parts containing a strong constitutive promoter, *TDH3p* (Figure 1). Even though the GoI was successfully integrated into all regions, the integrations efficiencies ranged from 35% to 41% (Figure 2C). Stratigopoulos et al. (2018) reported that 24 hours of DMSO feeding prior to CRISPR increased mammalian cell genome editing efficiency. ^63^ We, therefore, first tested whether DMSO feeding for 24 hours had a positive effect on CRISPR efficiency for *S. cerevisiae*. Unfortunately, DMSO feeding yielded inefficiently for integration rates with less than 30% (Figure 2C). Also, ∼50% fewer colonies were obtained after transformation compared to the DMSO-free method. Following that, the false positive colonies, which were able to grow on the selective media but did not express *mNeonGreen*, were further studied to determine whether they contained correctly assembled Cas9 plasmid, using primers flanking the overlapping sequences, connectors, as indicated in Figure 2A. This revealed that none of the false-positive colonies contained connector A, and only ∼10% of them had connector E in their plasmids. As both connectors were overlapping sequences flanking the *Cas9* cassette, we then tested if the *in vitro* plasmid parts could be amplified in full. The plasmid parts were used as a template, and ∼20 bp primers at the end of the terminals were used for a second PCR, as demonstrated in Figure 2B. Although the four plasmid parts, selection marker, yeast ORI, storage part, gRNA, were amplified, the Cas9 gene failed. In contrast, it was amplified from a correctly assembled plasmid, proving that the terminal sequences of the Cas9 gene (connector A and E) were not completely amplified by the Phusion polymerase. Indeed, there are limited studies about inefficient, repetitive PCR problems, which have contributed to the lack of reproducibility in certain assays in the synthetic biology community. ^64,65^ Shevchuk et al. (2004) reported the “shortening phenomenon” for PCR products. ^35^ Researchers revealed that the amplicons of high-fidelity DNA polymerases might yield truncated products because of the shortening of the amplicon’s ends, especially for relatively long targets (>3-4 kb). ^35^ For this reason, we applied two alternative approaches (i) using longer primers (∼120 bp ultramers, Table S1) containing non-functional extra bases at the terminal to nullify the shortening, (ii) dividing the long Cas9 gene with internal primers (Table S1) into two parts resulting ∼2.5 kb fragments. However, none of these strategies resulted in higher integration rates since integration rates similar to those of Phusion polymerase are achieved. Finally, we tested another high-fidelity DNA polymerase, PrimeSTAR GXL, which has a relatively higher error rate compared to Phusion. Surprisingly, it successfully amplified the Cas9 gene in full (Figure 2B), and all plasmid parts were re-amplified with a second PCR, as explained above. Consistently, the integration efficiencies in all regions increased dramatically by approximately 1.5 fold (*p-*value < 0.01), as shown in Figure 2C. Thus, all the plasmid parts were produced using PrimeSTAR GXL in the subsequent experiments, while donor parts were produced using Phusion polymerase.

**Figure 2:**
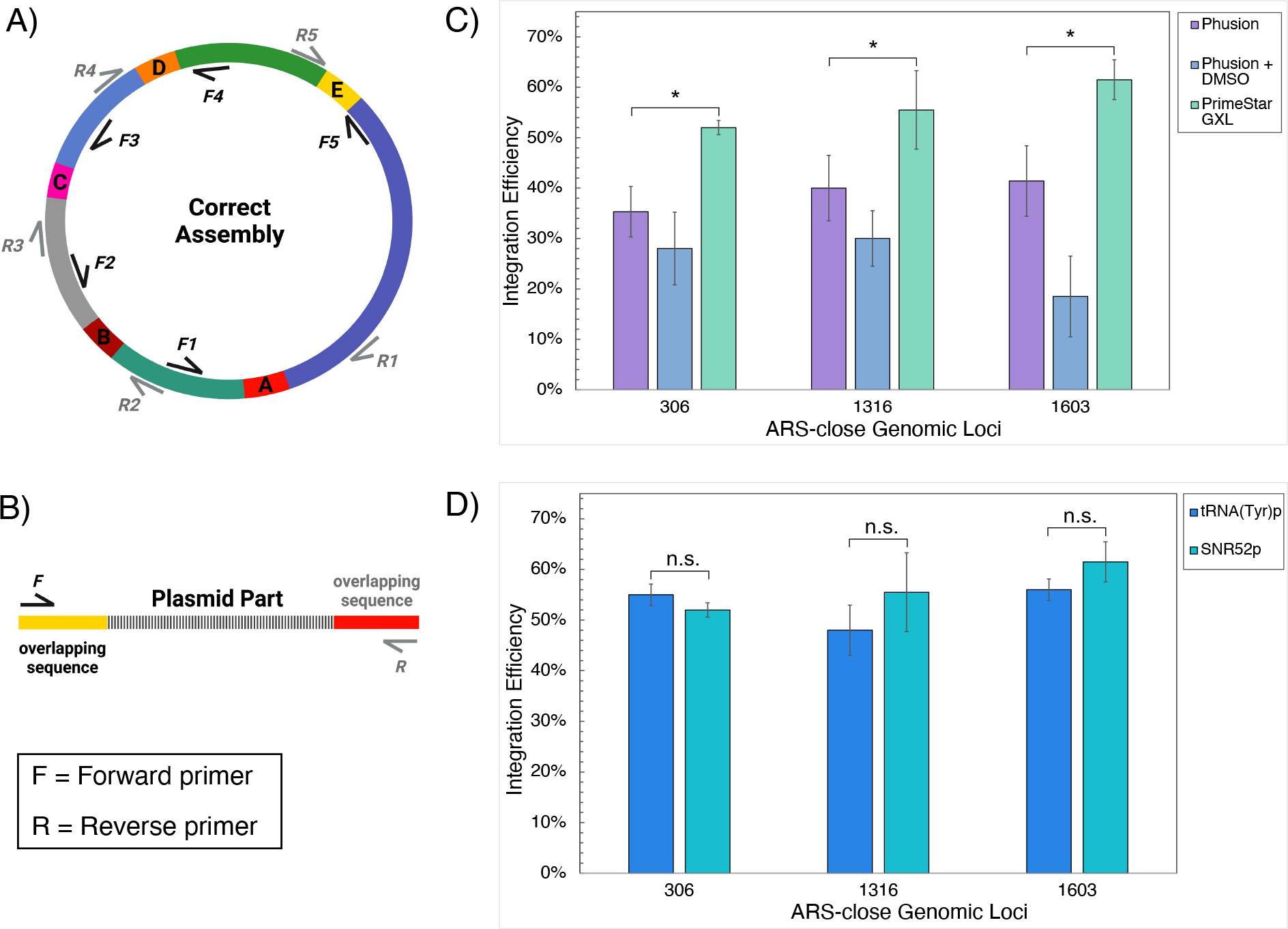
Optimization process and results of ACtivE. **A)** The all-in-one CRISPR plasmids were screened using plasmid colony PCR primers (F1-5, R1-5) targeting the connectors between adjacent plasmid modules. **B)** Each plasmid part was controlled to determine whether they were amplified in full after PCR reactions using ∼20 bp primers annealing at the end of the synthetic overlapping sequences. **C)** Comparison of integration efficiencies of different conditions; Phusion polymerase-based plasmid parts, DMSO feeding for 24 hours for Phusion polymerase-based plasmid parts, and PrimeSTAR GXL-based plasmid parts. **D)** Comparison of integration efficiencies when gRNA is expressed by two different promoters, *SNR52p* or *tRNA^(Tyr)^p*. The values are displayed as the mean of triplicate experiments, and error bars represent standard deviations. For simplification, ARS regions are shown with their corresponding numbers. The single asterisk represents a *p-*value < 0.01 and “n.s.” stands for not statistically significant (*p-*value > 0.05). The error bars show standard deviations of three replicates.

After improving the integration efficiency, false-positive colonies were screened to see if they contained the correctly assembled CRISPR plasmid after using the PrimeSTAR GXL. It was observed that ∼95% of the false-positive colonies had incorrectly assembled plasmids without gRNA and/or Cas9 parts, and all positive mNeonGreen expressing colonies had correct plasmids. This showed that the accuracy of *in vivo* plasmid assembly was the critical factor for CRISPR success in our method. The false-positive colonies also indicated that the plasmid could be assembled with missing parts. Different combinations of plasmid parts were transformed into yeast cells to confirm this. Although transformation yield dramatically decreased with missing parts, we observed a small number of colonies as long as they contained a selection marker and yeast ORI that are essential parts for surviving, as demonstrated in Figure S5. This was probably because of assembling the plasmid parts through non-homologous end-joining (NHEJ) rather than HDR. ^66,67^ NHEJ is a well-studied pathway, and the genes involved in this process are elucidated. ^66,67^ Therefore, as a further improvement, a disruption in the NHEJ pathway could increase the integration efficiency of ACtivE or similar methods, as shown previously for a non-conventional yeast *Yarrowia lipolytica*. ^68^

Additionally, we also compared two RNA expressing promoters to determine whether they affected gRNA expression and CRISPR activity as the differences in the promoters’ features might have an impact on the expression. ^37^ SNR52 RNA polymerase III promoter (*SNR52p*) is one of the most widely used promoters to express gRNAs in yeast. ^12,23^ Alternatively, gRNAs can be transcribed using tRNA promoters. ^11,40^ Therefore, we tested *SNR52p* and tRNA^(Try)^ promoters as shown in Figure 2D; however, no statistically significant difference (*p-*value > 0.05) was observed. Dong et al. (2020) compared tRNA^(Try)^, tRNA^(Pro)^, and *SNR52* promoters in terms of disruption yield on the *ADE2* gene and repression efficiency on a heterologous fluorescence protein. ^69^ Consistent with this study, researchers found very similar disruption and repression yields with these three gRNA promoters. ^69^ *SNR52p* consists of 269 bp, while *tRNA*^*(Try)*^*p* contains 118 bp (Figure S1). Therefore, promoters such as *tRNA*^*(Try)*^*p* could be used to minimize the cost of the synthetic gRNA cassettes.

### Integration Efficiencies on Eight ARS-close Genomic Loci

The improved ACtivE method was used to integrate *mNeonGreen* into five more ARS-close genomic loci to compare the integration efficiencies and characterize the selected eight genomic regions (Table 1). High-yield gene integrations ranging from 50% to 80% were achieved in the target regions, as shown in Figure 3A. Although 100% gene integration or deletion efficiencies have been previously reported,^12^ genome editing efficiency of ACtivE is remarkably high considering its convenience and rapidity. In addition, previous works reported extremely low (< 20%) CRISPR-based heterologous gene integration efficiencies in some genomic locations. ^40^ Therefore, the regions tested in this study are suitable targets for gene integrations.

**Figure 3:**
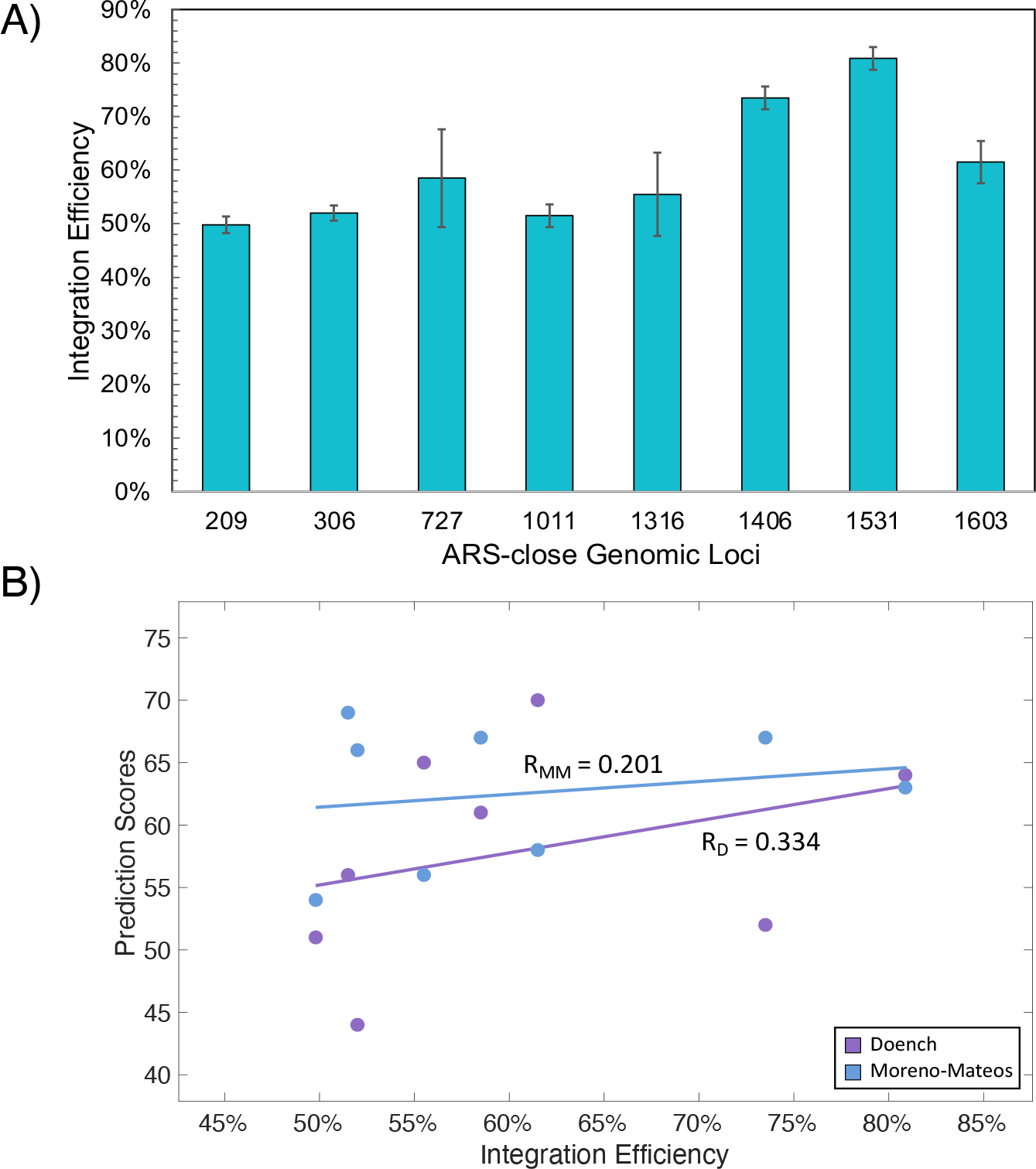
Integration efficiencies and their correlation with prediction algorithms. **A)** Integration efficiency of *mNeonGreen* on eight ARS-close genomic loci. The values are displayed as the mean of triplicate experiments and error bars represent standard deviations. For simplification, ARS regions are shown with their corresponding numbers. **B)** Linear regression models showing the correlation between integration percentages and gRNA efficiency scores of Doench’s ^49^ and Moreno-Mateos’ ^50^ prediction algorithms. R_D_ = Pearson correlation coefficient based on Doench’s algorithm, R_MM_ = Pearson correlation coefficient based on Moreno-Mateos’ algorithm. The error bars show standard deviations of three replicates.

The relationship between integration efficiencies and gRNA efficiency scores of the genomic regions (Table 1) was also determined using linear regression models, as shown in Figure 3B. Moderate and weak positive correlations were found on Doench ^49^ and Moreno-Mateos ^50^ algorithms with a Pearson’s correlation coefficient of R=0.334 and R=0.201, respectively. These findings evaluating a small dataset can be considered promising, although the best fitting gave only a moderate positive correlation. These findings suggest that computational gRNA design/scoring tools can help with selecting gRNA sequences for CRISPR-based studies in yeast, considering the algorithms’ limitations. Besides this, a larger sample size is required to properly evaluate these prediction algorithms, which should be further improved by considering, for example, 3-D chromatin structures of the yeast ^70^ to develop more reliable, yeast-specific prediction tools.

### Single-step Gene Deletion Using Only Homology Arms

Besides genomic integration, gene deletion was also tested using ACtivE. To this end, two non-essential genes, the *GAL80* gene encoding a regulator protein for galactose-related metabolic genes ^71^ and the *DIT1* encoding a sporulation-specific enzyme,^72^ were deleted. Using only UHA and DHA flanking outside the genes (primers and crRNAs are listed in Table S5), ∼75% deletion efficiency (standard deviation ≈ 8%) and ∼93% deletion efficiency (standard deviation ≈ 11%) were achieved for the *GAL80* and *DIT1* genes, respectively, without the need for any heterologous gene part. In this way, the whole *GAL80* and *DIT1* genes were deleted without any scar. The deletions were confirmed using colony PCR and Sanger sequencing. This study demonstrates the flexibility of ACtivE as the same approach can be also used for scar-free gene deletion.

### Characterization of Genomic Loci

The genomic regions used as landing pads for *mNeonGreen* were characterized in terms of gene expression rate and effect on the cell fitness in two different media, YPD and SD, as described in the materials and methods section. First, plasmid-free cells were selected using 5-FOA counter-selection to eliminate plasmid burden. OD_600_ was measured to compare biomass and growth rates of the strains containing mNeonGreen on different genomic loci. Figure 4 shows the biomass of the yeast strains in YPD and SD media.

**Figure 4:**
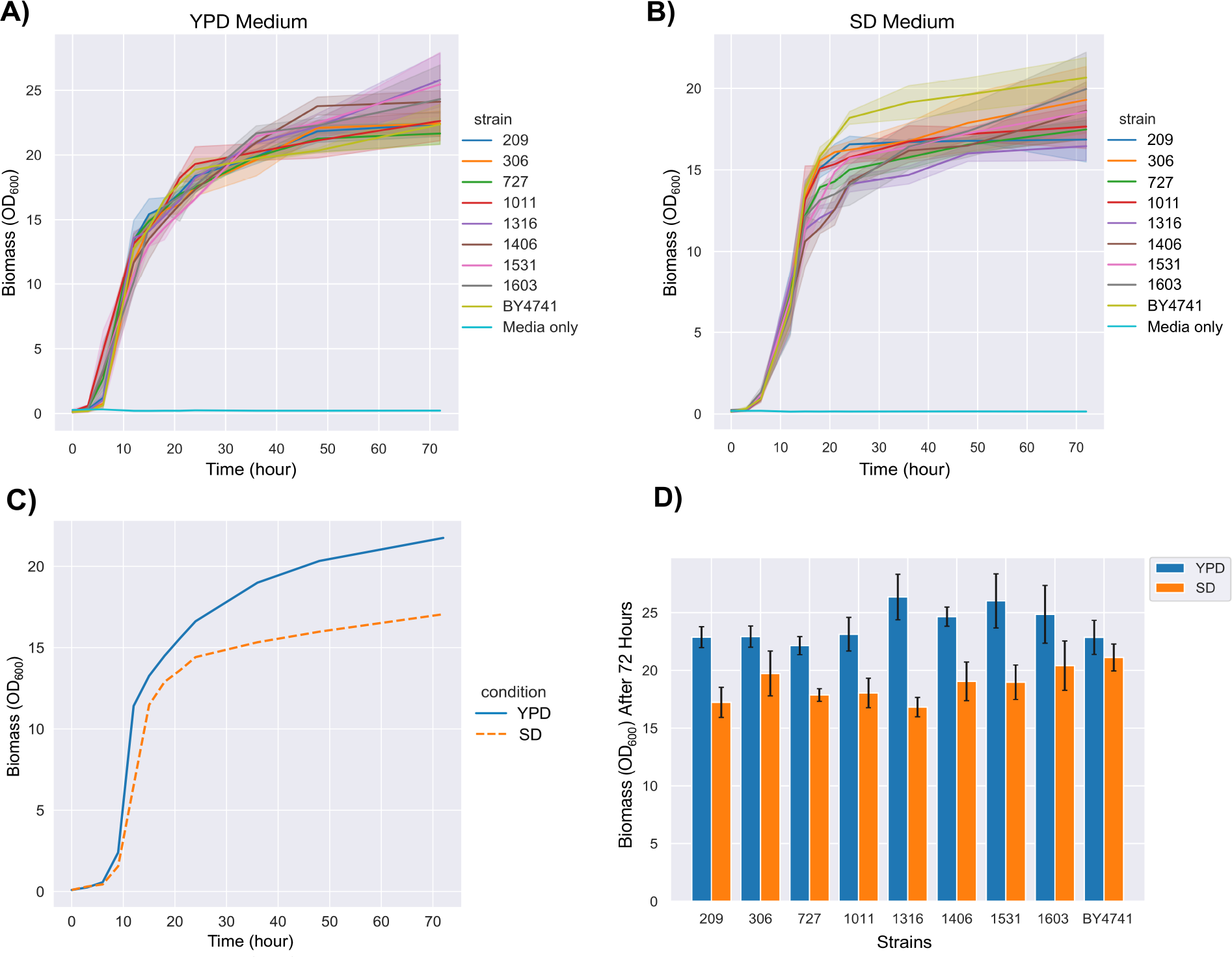
The biomass comparison between *mNeonGreen* integrated strains and the parental strain **A)** The biomass over 72 hours in YPD media **B)** The biomass over 72 hours in SD media **C)** Growth trends in YPD and SD media comparing mean biomass of all strains in these two conditions over 72 hours **D)** The final OD_600_ values represent the total biomass after 72 hours. The solid lines on the curves stand for the average values of independent colonies or different strains. The standard deviations of the three replicates are shown by shading on the curves or by error bars on the bar chart.

As seen, the growth curves presented the expected trends in both media (Figure 4A & 4B). The average biomass (Figure 4C) of nine strains, including the parental strain, BY4741, was expectedly higher in YPD media compared to SD media (*p-*value < 0.01) as YPD is a richer environment than SD media.^73,74^ The biomass order in YPD media was as follows:

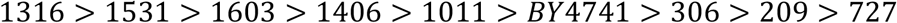

Although the 1316 (OD_600_ *≈* 26) showed the highest biomass, and the 727 (OD_600_ *≈* 22) showed the lowest biomass at the 72^nd^ hour (Figure 4A), there were no statistically significant differences when they were compared to the parental strain (OD_600_ *≈* 22.5) (*p-*value > 0.05). This data showed that perturbations on these genomic loci are unlikely to cause a negative effect on cell fitness considering the parental strain. Apel et al. (2017) reported a similar result as they did not find a significant difference in growth rates of GFP expressing strains compared to their parental strain in YPD media. ^40^ However, a statistically significant difference (*p-*value < 0.05) was observed between the 1316 and the 727, showing that targeting ARS1316 likely yields better biomass than ARS727 in YPD media.

On the other hand, the average biomass order in SD media was as follows:

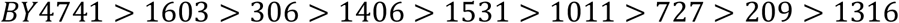

BY4741 showed the highest biomass amount as the parental strain in SD media (Figure 4B). Besides, a statistically significant difference (*p-*value < 0.05) was observed when the parental strain BY4741 was individually compared with 209, 727, 1011, and 1316, meaning that genetic perturbations on these loci might negatively affect the cell fitness in SD media. Surprisingly, 1316 resulted in the lowest biomass in SD media even though it was the best growing strain in YPD. Also, a substantially different biomass order was observed in SD media compared to YPD media (Figure 4D). These results show that genetic alterations in the corresponding locations might distinctly affect the cell fitness of yeast strains depending on the environmental conditions and the media compositions.

In addition, the growth rates of the strains were calculated using a Gaussian process-based algorithm. ^75^ As shown in Figure S6, the maximum growth rate of BY4741 was observed around the 6^th^ hour in both YPD and SD media. Similarly, the maximum growth rates of *mNeonGreen* integrated strains were at the 6^th^ hour in SD media (Figure S8). However, 1011, 1406, 1531, and 1603 showed the maximum growth rate at around the 3^rd^ hour, whereas the others were around the 6^th^ hour (Figure S7). Probably, perturbations on ARS1011, ARS 1406, ARS1531, and ARS1603 affect the growth rate in YPD media.

To compare gene expression, fluorescence intensities produced by mNeonGreen were measured. Autofluorescence caused by the yeast cell itself and the media ^46^ was corrected using *omniplate* ^53^ to detect the fluorescence intensity per OD.

All strains showed similar expression patterns based on the selected medium, YPD or SD, with some fluctuations in the first 24 hours, as shown in Figures 5A & 5B. As mNeonGreen expression was driven by the constitutive *TDH3* promoter in all mNeonGreen expression strains, these results also showed the expression patterns of *TDH3p* in YPD and SD media (Figure 5C). In YPD media, the best mNeonGreen expressing strains depended on the hour. For instance, at 24^th^, 36^th,^ and 72^nd^ hours, the 209 was the best strain for mNeonGreen expression, whereas the 727 showed the highest expression at the 48^th^ hour (Figure 5A). These two strains were also the most mNeonGreen producing strains in 72 hours (Figure 5D).

**Figure 5:**
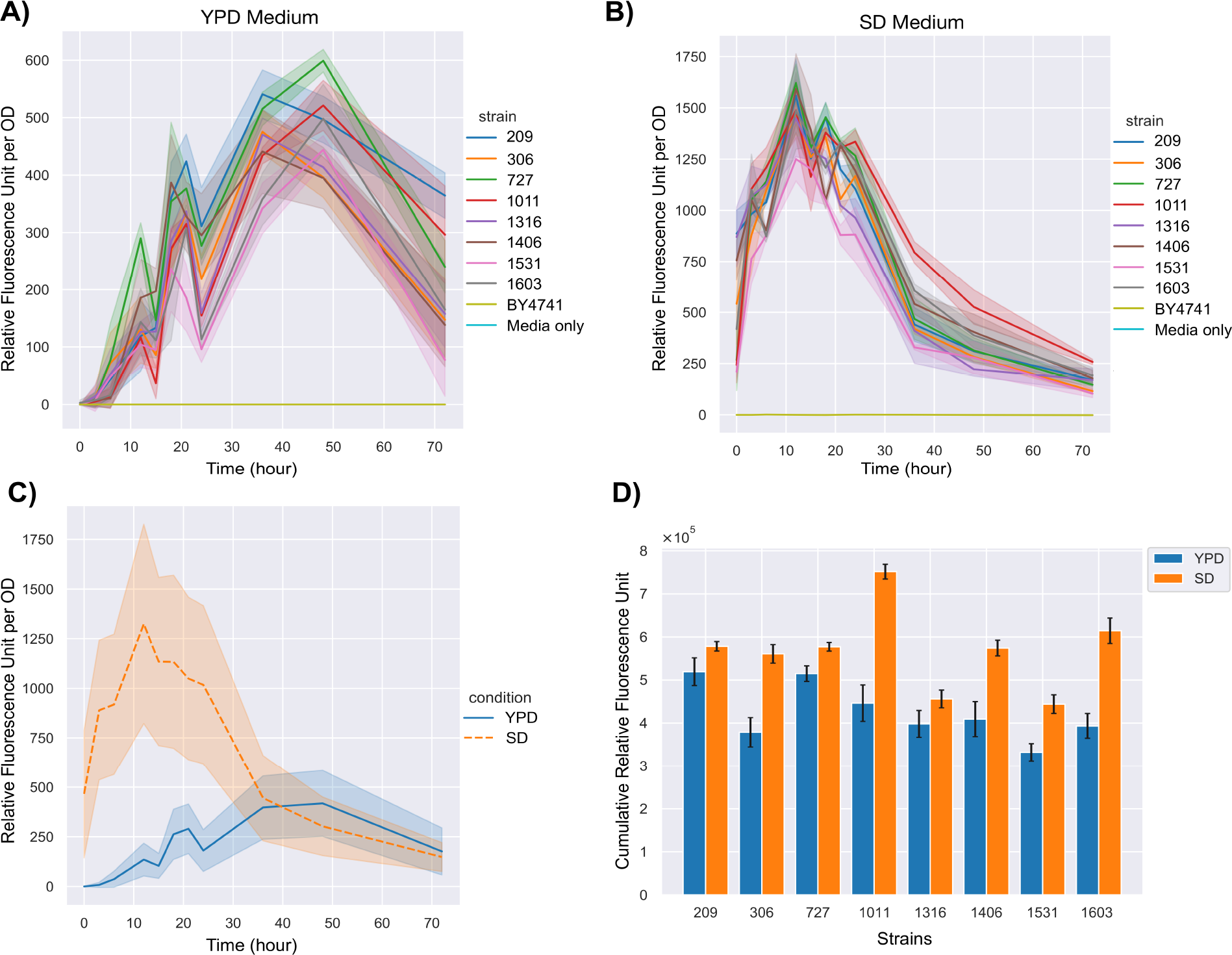
The comparison of heterologous gene expression between *mNeonGreen* integrated strains **A)** Relative fluorescence intensities (RFU) over 72 hours in YPD media **B)** RFUs over 72 hours in SD media **C)** The average RFUs of all strains in YPD and SD media separately. **D)** The cumulative RFU values represent the total gene expression or protein production by the total biomass at each time for 72 hours. The standard deviations of the three replicates are shown by shading on the curves or by error bars on the bar chart. The solid lines on the curves stand for the average values of independent colonies or different strains.

On the other hand, 1011 showed the highest expression from the 24^th^ hour in SD media (Figure 5B). Therefore, the ARS1011 was the best integration site for the cumulative expression of mNeonGreen in SD media even though the 1011’s biomass was significantly lower than the parental strain (*p-*value < 0.05) in SD media (Figure 4D). Moreover, Figures 4C & 5C show that more biomass was obtained in YPD media. Still, mNeonGreen expression was dramatically higher in SD media as there was a significant difference in expression rates in the first 36 hours of SD and YPD media. This shows that the expression rates were maximum in the exponential phase in SD media. However, after the 24th hour, there was a sharp decrease in the fluorescence unit. This decrease might be because of both reduction in the expression and the degradation of the mNeonGreen. Nevertheless, the cumulative expression indicating the total protein production over 72 hours by total biomass at each time can be helpful information, especially for protein secretion into the extracellular environment where the effect of intracellular degradation is minimum. Therefore, SD media was more advantageous for cumulative protein production than YPD media for these genomic loci.

The time-derivatives of mNeonGreen expressions were also found, as shown in Figures S9 & S10. The accelerations in expression reached the maximum between 10^th^ and 20^th^ hours in YPD media except for 1531, which showed the maximum increase in the expression rate between 30^th^ and 40^th^ hours (Figure S9). In contrast, the increase in the expression rates was the highest in the first 10 hours in SD media for all mNeonGreen expressing strains (Figure S10). These results were consistent with Figure 5C as the expression started with ∼500 relative fluorescence units (RFU) in SD media and reached the maximum at around the 12^th^ hour, while it began with ∼0 RFU and reached the maximum at around the 48^th^ hour in YPD media (Figure 5C).

Indeed, locus-based, *TDH3p-*driven heterologous gene expression in the yeast genome has been thoroughly studied in this work, presenting biomass, fluorescence intensity per OD, expression rate, and cumulative expression amount in 72 hours with 11 different time points in two different media. In a similar study, Apel et al. (2017) compared the GFP fluorescence levels among 23 genomic loci at the 8^th^ and 24^th^ hours in YPD and SD media using a *TEF1p promoter* instead of the *TDH3p* used in this work. ^40^ The researchers reported a higher average fluorescence level in YPD at the 8^th^ hour, whereas it was clearly in favor of SD medium at the 24^th^ hour. ^40^ In the same study, the promoters were also compared, and it was shown that the fluorescence level was the highest in SD media with *TDH3p-*driven expression at the 4^th^ hour on a single locus, whereas it was slightly higher at the 8^th^ and 24^th^ hours in YPD. ^40^ Although having a higher fluorescence intensity in SD media with *TDH3p* in the early hours of the cultivation overlaps with our results; we observed a clear difference in fluorescence intensity in favor of SD media in the first 32 hours (Figure 5C) that was longer than it was reported in the previously mentioned reference. ^40^ These findings can also give insights into the expression patterns of the native *TDH3* gene that encodes an enzyme, glyceraldehyde 3-phosphate dehydrogenase, involved in glycolysis, transcriptional silencing, and rDNA recombination. ^76^

### Multiplexing using ACtivE

Simultaneous genome alterations in a single step can be preferential to accelerate genome editing or heterologous pathway construction. Therefore, the multiplexing capacity of ACtivE was tested for multi loci and multi-gene integrations into the yeast genome using different plasmid assembly and donor DNA delivery strategies, as illustrated in Figure 6. Initially, two fluorescent reporter proteins, mNeonGreen and mCherry, were integrated into ARS1406 and ARS1531 loci, respectively (Figure 7B), using two independent gRNA modules to be assembled into two different plasmids (Figure 6A). With this strategy, *mNeonGreen* and *mCherry* were simultaneously integrated into 21% of the colonies (Figure 7C). Alternatively, the gRNAs targeting ARS1406 and ARS1531 were *in vivo* assembled using synthetic homology sequences resulting in a single all-in-one plasmid expressing both gRNAs simultaneously (Figure 6B). The integration rate reached 46% with this approach (Figure 7C).

**Figure 6:**
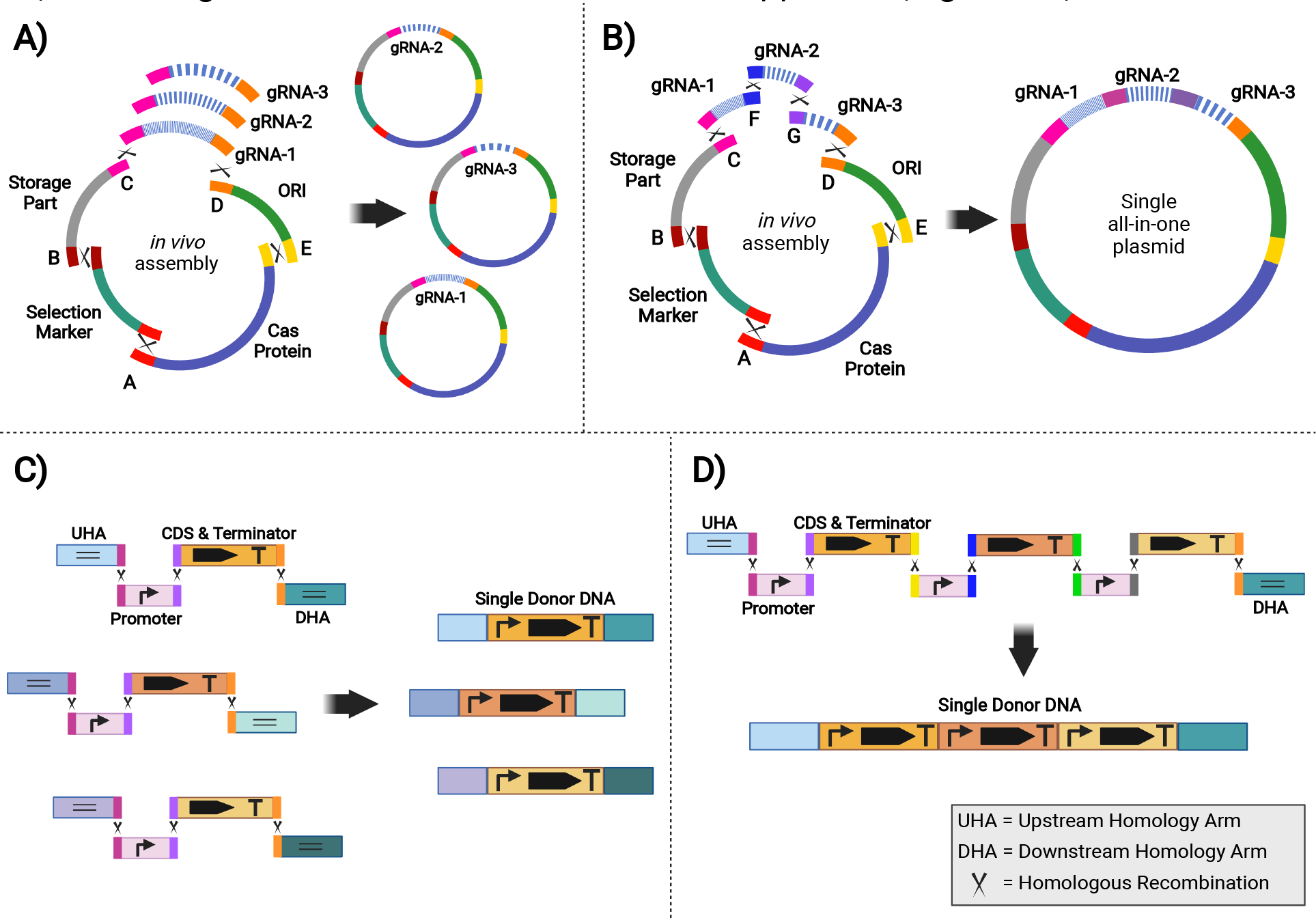
Alternative approaches for multiplexing **A)** Different gRNA cassettes are *in vivo* assembled, resulting in different plasmids targeting multiple loci. Each gRNA cassette contains identical overlapping sequences at their terminals so that each one individually assembles with the other plasmid parts. **B)** The gRNA cassettes are tandemly assembled, resulting in a single all-in-one plasmid that targets multiple loci. **C)** For multi-locus and multigene integration, each gene has its own homology arms (HA) depending on the target site. **D)** The promoters and CDSs can be tandemly assembled for single-locus multi-gene integration. The multi-gene cassette contains a single upstream homology arm (UHA) and downstream homology arm (DHA).

**Figure 7:**
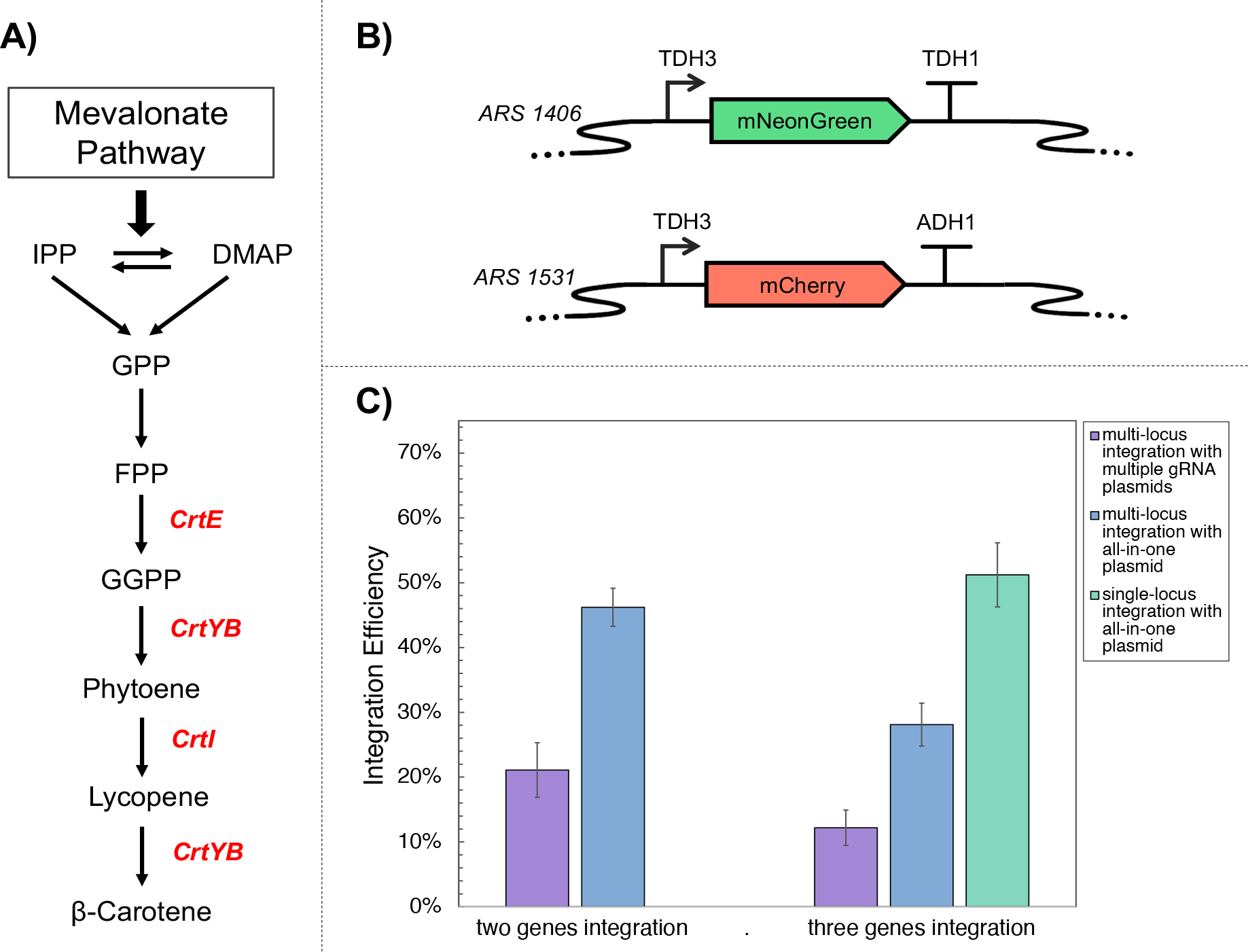
**A)** The heterologous β-carotene pathway constructed in a single-step multi-gene integration. The heterologous genes are shown in red. **B)** Genetic construct illustrations of *mNeonGreen* and *mCherry* genes used for single-step double gene integration and *CrtE, CrtI*, and *CrtYB* genes used for single-step triple gene integration. **C)** Integration efficiencies achieved using different strategies. The single asterisk represents a *p-*value < 0.01. The error bars show standard deviations of three replicates.

Following this approach, three heterologous genes of the β-carotene pathway, *CrtE, CrtYB, and CrtI* (Figure 7A), were simultaneously integrated into ARS1406, ARS1531, and ARS1603, respectively. A 12 % integration rate was achieved using the strategy of assembling independent plasmids for each gRNA (Figure 6A), but this increased to 28% when three gRNAs were assembled in a single plasmid (Figure 6B). Furthermore, the genes were integrated into a single locus (ARS1531) to construct the multi-gene pathway on a single genomic location (Figure 6D), and 51% of the colonies successfully produced the orange pigment. Indeed, the differences in integration efficiencies were not surprising as higher integration rates were observed with fewer linear fragments. Even though the 12% integration rate is relatively low, the correct assembly and/or integration of 19 linear fragments (Figure 6A & 6C) shows the flexibility and capability of the ACtivE method.

## CONCLUSION

The ACtivE method proposes a practical strategy to accelerate yeast genome engineering. It eliminates the use of any reagents or kits used for *in vitro* DNA assembly and bypasses extra cloning steps. Therefore, verified linear fragments have been collected to create a flexible CRISPR toolkit for many different purposes. The first version of the toolkit and a user manual are freely available at https://www.leorioslab.org/cost-crispr-toolkit/. The user can simply combine the modules provided in the toolkit without any plasmid purification, enzymatic treatment, *in vitro* assembly, or cloning steps. Providing custom donor DNA, the whole work for genome editing can be completed in only one day using verified linear fragments. Moreover, the modules to be used for CRISPR plasmid construction can be stored for a long time for further applications, and they can be exchanged between different groups as standard parts thanks to their synthetic ends. In this way, customized CRISPR plasmids containing specific parts such as Cas protein, selective marker, gRNA, yeast ORI or bacterial marker can be easily obtained.

To increase the genome editing efficiency and reproducibility of this method, the importance of the DNA polymerase type to be used for part amplification was underlined. In other words, the plasmid modules had to be completely amplified as their ends are critical for *in vivo* assembly. To this end, an appropriate DNA polymerase synthesizing complete amplicons containing the connectors was used to overcome the shortening problem of the DNA fragments that might be a bottleneck to amplify long amplicons.

ACtivE achieved more than 80% integration for a single gene in the ARS1531 region. For multiplexing, expressing multiple gRNAs through a single plasmid resulted in a higher integration yield for multi-loci and multi-gene integration. More than 50% integration efficiency for triple genes was reached in the ARS1531 region as a single-locus multi-gene integration. Also, more than 90% deletion efficiency was obtained on the *DIT1* gene. Considering its usefulness and pace, this method should accelerate genome editing processes as the strains of interest can be smoothly detected after a simple screening.

In addition, eight ARS-close regions in the yeast genome were thoroughly characterized using two different yeast media. The total biomass and growth rates of the strains containing heterologous genes in the corresponding loci were found. Also, RFUs were detected to characterize the gene expression rates and total protein production in these locations. All strains showed higher growth rates in YPD than in the SD medium. Considering the growth rates, ARS 1316 and ARS 1531 might be good targets for high biomass in YPD, whereas ARS 306 and ARS 1603 are preferable for higher biomass in SD media. Nonetheless, no significant difference in the final biomass was observed compared to the parental strain in YPD media. In contrast, strains 209, 727, 1011, and 1316 showed lower growth rates than the parental strain in SD media. On the other hand, gene expression rates might vary depending on the locus used. ARS 209 and ARS 727 showed better protein expressions in YPD media, while ARS 1011 was the best locus for gene expression in SD media. Besides, dynamic gene expression varied considerably mainly depending on the medium conditions, and the cumulative expressions were higher in SD media, although the biomasses were lower in this media.

*S. cerevisiae* is an important chassis organism for many applications, from metabolic engineering to disease modeling. The CRISPR/Cas system has been a versatile instrument for designing its genome. The improvements and alternative approaches presented in this paper have a great potential to accelerate the yeast genome editing process in a standardizable and easy way. The genomic loci characterized in this study provide more options for well-defined genomic landing sites, especially for yeast cell factory design.

## Notes

The authors declare no competing financial interest.

## Data Availability

The detailed information about the parts and modules in the toolkit and the version updates can be found at https://www.leorioslab.org/cost-crispr-toolkit/. The codes and the standard curve for plate reader data analysis can be found on https://github.com/kmalci/plate-reader/.

## Acknowledgments

This study was supported by the YLSY Program of the Ministry of National Education of the Republic of Turkey, the British Council (Grant Number: 527429894), IBCatalyst (Project No. 102297), Biotechnology and Biological Sciences Research Council (BBSRC; Project BB/S017712/1). The authors thank Dr. Dariusz Abramczyk for providing the BY4741 strain, Dr. Matthew Dale for his gift of vector pCPS1ULA-BA6, Dr. Jamie Auxillos for her gift of pTDH3-Re2.8-2 vector, Alexander Speakman for technical assistance about the microplate reader, and Prof. Peter Swain for his guide on the *omniplate* software.

