## Supplemental File for "ACtivE: Assembly and CRISPR-targeted *in vivo* Editing for Yeast Genome Engineering Using Minimum Reagents and Time"

**Table S1:** The primer list for *in vivo* CRISPR plasmid assembly and for controlling correct assembly

| Name | Sequence 5' to 3' | Purpose |
|---|---|---|
| yCas9 + E For | CATACGTTGAAACTACGGCAAAGGATTGGTCAGATCGCTTCATACAGGGAAAGTTCGGCAGAATTCGCATCTAGACTGAACTG | yCas9 amplification with synthetic overlapping sequences |
| yCas9 + A Rev | GTGCCTATTGATGATCTGGCGGAATGTCTGCCGTGCCATAGCCATGCCTTCACATATAGTGAGGTAGGGCATATGTCCTCTG | |
| yCas9 + E (ultramer) For | TGTCATACAGCTCAGGGATTGGTCAAGGATTCTTCATACACATACGTTGAAACTACGGCAAAGGATTGGTCAGATCGCTTCATACAGGGAAAGTTCGGCAGAATTCGCATCTAGACTGAACTG | yCas9 amplification with synthetic overlapping sequences and extra bases |
| yCas9 + A (ultramer) Rev | AGCCATAGCGAATGTCTGTCTGCATTATAGGTCTGCCGTGGTGCCTATTGATGATCTGGCGGAATGTCTGCCGTGCCATAGCCATGCCTTCACATATAGTGAGGTAGGGCATATGTCCTCTG | |
| yCas9 Internal For | GACATTGTCCTGACTCTCACTCTGTTCGAGGACCGGGAAATGATCGAGGAG | Amplification of half-length yCas9 |
| yCas9 Internal Rev | CTTAAGCCTCTCCTCGATCATTTCCCGGTCCTCGAACAGAGTGAGAGTCAG | |
| URA3 + A For | ACTATATGTGAAGGCATGGCTATGGCACGGCAGACATTCCGCCAGATCATCAATAGGCACGCAGATTGTACTGAGAGTGCACC | URA3 amplification with synthetic overlapping sequences |

| | | |
|---|---|---|
| URA3 + B Rev | GTTGAACATTCTTAGGCTGGTCGAATCATTTAGACACGGGCATCGTCCTCTCGAAAGGTGCGCATCTGTGCGGTATTTCAC | |
| *E.coli* Part + B For | CACCTTTCGAGAGGACGATGCCCGTGTCTAAATGATTCGACCAGCCTAAGAATGTTCAACGTGCGCGGAACCCCTATTTG | Amplification of bacterial fragment with synthetic overlapping sequences |
| *E.coli* Part + C Rev | CTAGCGTGTCCTCGCATAGTTCTTAGATTGTCGCTACGGCATATACGATCCGTGAGACGTGAGCGGTATCAGCTCACTCAAAG | |
| gRNA cassette + C For | ACGTCTCACGGATCGTATATGCCGTAGCGACAATCTAAGAACTATGCGAGGACACGCTAGCCCTCACTAAAGGGAACAAAAGC | Amplification of gRNA cassette with synthetic overlapping sequences |
| gRNA cassette + D Rev | AATCACTCTCCATACAGGGTTTCATACATTTCTCCACGGGACCCACAGTCGTAGATGCGTGGAAACAACAAAAGGATGTCGAC | |
| gRNA cassette For | ACGTCTCACGGATCGTATATG | Amplification of gRNA cassette for tandem gRNAs for multiplexing |
| gRNA cassette + F Rev | AAGGGCCATGACCACCTGATGCACCAATTAGGTAGGTCTGGCTATGTCTATACCTCTGGCTAAATTGGCCATAGAAAAATTCTGTTATC | |
| gRNA cassette + F For | GCCAGAGGTATAGACATAGCCAGACCTACCTAATTGGTGCATCAGGTGGTCATGGCCCTTTGGAGCTCTTTGAAAAGATAATGTATG | Amplification of gRNA cassette for tandem gRNAs for multiplexing |
| gRNA cassette + G Rev | GTCACGGGTTCTCAGCAATTCGAGCTATTACCGATGATGGCTGAGGCGTTAGAGTAATCTTAAATTGGCCATAGAAAAATTCTGTTATC | |
| gRNA cassette + G For | AGATTACTCTAACGCCTCAGCCATCATCGGTAATAGCTCGAATTGCTGAGAACCCGTGACTGGAGCTCTTTGAAAAGATAATGTATG | Amplification of gRNA cassette for tandem gRNAs for multiplexing |
| gRNA cassette Rev | CAATCACTCTCCATACAGGGTTTC | |
| 2μ ori + D For | ACGCATCTACGACTGTGGGTCCCGTGGAGAAATGTATGAAACCCTGTATGGAGAGTGATTGACGAAAGGGCCTCGTGATAC | Amplification of 2μ ori with synthetic overlapping sequences |
| 2μ ori + E Rev | TGCCGAACTTTCCCTGTATGAAGCGATCTGACCAATCCTTTGCCGTAGTTTCAACGTATGCATTTCCCCGAAAAGTGCCACC | |
| A For | GGAAGTCAAGAAGGACCTTATC | Colony PCR for fragment A |
| A Rev | GTGTGCATTCGTAATGTCTG | |
| B For | CGGCAGAAGAAGTAACAAAG | Colony PCR for fragment B |
| B Rev | GTCATGCCATCCGTAAGATG | |
| C For | CGAACTGAGATACCTACAGC | Colony PCR for fragment C |
| C Rev | CAAGTTGATAACGGACTAGCC | |
| D For | CTATTGTTATGTAAAATGCCACCT | Colony PCR for fragment D |
| D Rev | CACATACAGCTCACTGTTCA | |
| E For | GAAGCACAGATTCTTCGTTGG | Colony PCR for fragment E |
| E Rev | GTTCCTCACTCTTTCCTTACTCA | |

* yCas9 stands for yeast codon-optimized Cas9 gene

** Black nucleotides represent the annealing part; red nucleotides represent the overlapping part of the primers. Blue nucleotides in the ultramers were used to lengthen the primers, and they do not have any other function.

*** A, B, C, D, and E stand for 60 bp synthetic overlapping fragments

**Table S2:** The connectors (60 bp synthetic fragments) used for *in vivo* plasmid assembly

| ***Name*** | ***Sequence 5' to 3'*** |
|---|---|
| ***A*** | ACTATATGTGAAGGCATGGCTATGGCACGGCAGACATTCCGCCAGATCATCAATAGGCAC |
| ***B*** | CACCTTTCGAGAGGACGATGCCCGTGTCTAAATGATTCGACCAGCCTAAGAATGTTCAAC |
| ***C*** | ACGTCTCACGGATCGTATATGCCGTAGCGACAATCTAAGAACTATGCGAGGACACGCTAG |
| ***D*** | ACGCATCTACGACTGTGGGTCCCGTGGAGAAATGTATGAAACCCTGTATGGAGAGTGATT |
| ***E*** | CATACGTTGAAACTACGGCAAAGGATTGGTCAGATCGCTTCATACAGGGAAAGTTCGGCA |
| ***F*** | GCCAGAGGTATAGACATAGCCAGACCTACCTAATTGGTGCATCAGGTGGTCATGGCCCTT |
| ***G*** | AGATTACTCTAACGCCTCAGCCATCATCGGTAATAGCTCGAATTGCTGAGAACCCGTGAC |

* The fragments are orthogonal with 46% < GC < 51% [1]

**Table S3:** The primer list for *mNeonGreen* integration into eight genomic loci and *mCherry* integration into ARS 1531 region

| **Name** | **Sequence 5' to 3'** | **Purpose** |
|---|---|---|
| 209 UHA For | GCAAAGAGCAATGGCAACAG | Amplification of 209 UHA with an overlapping sequence for *TDH3p* |
| 209 UHA Rev | TATTCTTTGAAATGGCAGTATTGATAATGAC<br>TAGCACATTTTATGGGCCTAAG | |
| 209 *TDH3p* For | ATGATGTCTTAGGCCCATAAAATGTGCTAG<br>TCATTATCAATACTGCCATTTCAAAG | Amplification of promoter *TDH3p* with overlapping sequences for 209 UHA and *mNeonGreen* |
| *TDH3p* Rev (**common for all regions**) | CATGTTATCCTCCTCGCCCTTGCTCACCAT<br>TTTGTTTGTTTATGTGTGTTTATTCGAAAC | |
| *mNeonGreen* For (**common for all regions**) | GTTTCGAATAAACACACATAAACAAACAAA<br>ATGGTGAGCAAGGGCGAGGA | Amplification of *mNeonGreen* with overlapping sequences for *TDH3p* and 209 DHA |
| 209 *mNeonGreen* Rev | TTGAATACAGAGCAAAAGGATTAGCCATAC<br>CGTTCAGGGTAATATATTTTAACCGCCG | |
| 209 DHA For | GTCGGCGGTTAAAATATATTACCCTGAACG<br>GTATGGCTAATCCTTTTGCTCTG | Amplification of 209 DHA with an overlapping sequence for *mNeonGreen* |
| 209 DHA Rev | CTCTATATCGCTGTTGCTTATGG | |
| 306 UHA For | GTGACTGTCTCCAAGAATACGAC | Amplification of 306 UHA with an overlapping sequence for *TDH3p* |
| 306 UHA Rev | TATTCTTTGAAATGGCAGTATTGATAATGAC<br>GTTATTGATGTTAGGAGAAGGAGC | |
| 306 *TDH3p* For | CTGTTCGTTATTGATGTTAGGAGAAGGAGC<br>TCATTATCAATACTGCCATTTCAAAG | Amplification of promoter *TDH3p* with overlapping sequences for 306 UHA and *mNeonGreen* |

| | | |
|---|---|---|
| 306 *mNeonGreen* Rev | TTCAGAAACACTGCTTACACTATTCACCAG<br>CGTTCAGGGTAATATATTTTAACCGCCG | Amplification of *mNeonGreen* with overlapping sequences for *TDH3p* and 306 DHA |
| 306 DHA For | GTCGGCGGTTAAAATATATTACCCTGAACG<br>CTGGTGAATAGTGTAAGCAGTGTTTC | Amplification of 306 DHA with an overlapping sequence for *mNeonGreen* |
| 306 DHA Rev | CAAGAACACCAGACCTCCAAGC | |
| 727 UHA For | CTGCCCCAGGACTTGGAAGG | Amplification of 727 UHA with an overlapping sequence for *TDH3p* |
| 727 UHA Rev | TATTCTTTGAAATGGCAGTATTGATAATGAC<br>ATAGCAGTGGCGCGGTC | |
| 727 *TDH3p* For | TTATAGGGAATCGACCGCGCCACTGCTAT<br>GTCATTATCAATACTGCCATTTCAAAG | Amplification of promoter *TDH3p* with overlapping sequences for 727 UHA and *mNeonGreen* |
| 727 *mNeonGreen* Rev | ATCAGCAGGCCATGGATAAACTTTCCGTTG<br>CGTTCAGGGTAATATATTTTAACCGCCG | Amplification of *mNeonGreen* with overlapping sequences for *TDH3p* and 727 DHA |
| 727 DHA For | GTCGGCGGTTAAAATATATTACCCTGAACG<br>CAACGGAAAGTTTATCCATGG | Amplification of 727 DHA with an overlapping sequence for *mNeonGreen* |
| 727 DHA Rev | GAGATTCTTGACGTAAAGTGC | |
| 1011 UHA For | GTGGTACAAGAAGCGTTGGAGAC | Amplification of 1011 UHA with an overlapping sequence for *TDH3p* |
| 1011 UHA Rev | TATTCTTTGAAATGGCAGTATTGATAATGAC<br>TTCCAGCAGCGCCAGTAG | |
| 1011 *TDH3p* For | GCTCAACAACCCTACTGGCGCTGCTGGAA<br>GTCATTATCAATACTGCCATTTCAAAG | Amplification of promoter *TDH3p* with overlapping sequences for 1011 UHA and *mNeonGreen* |
| 1011 *mNeonGreen* Rev | CCAACACTTGATAGTATCTACTCGCCATTC<br>CGTTCAGGGTAATATATTTTAACCGCCG | Amplification of *mNeonGreen* with overlapping sequences for *TDH3p* and 1011 DHA |
| 1011 DHA For | GTCGGCGGTTAAAATATATTACCCTGAACG<br>GAATGGCGAGTAGATACTATCAAG | Amplification of 1011 DHA with an overlapping sequence for *mNeonGreen* |
| 1011 DHA Rev | CTTCACATTGAGTTTGAATATGCC | |
| 1316 UHA For | GGTTTCAAGCCAAATTGTACG | Amplification of 1316 UHA with an overlapping sequence for *TDH3p* |
| 1316 UHA Rev | TATTCTTTGAAATGGCAGTATTGATAATGAC<br>TTAGGTAGTAACTATACGCAGC | |
| 1316 *TDH3p* For | GGAGCAGCTGCGTATAGTTACTACCTAAGT<br>CATTATCAATACTGCCATTTCAAAG | Amplification of promoter *TDH3p* with overlapping sequences for 1316 UHA and *mNeonGreen* |
| 1316 *mNeonGreen* Rev | TAGCCCACTTCTAGCCAACTTCTAGCCCAC<br>CGTTCAGGGTAATATATTTTAACCGCCG | Amplification of *mNeonGreen* with overlapping sequences for *TDH3p* and 1316 DHA |
| 1316 DHA For | GTCGGCGGTTAAAATATATTACCCTGAACG<br>GTGGGCTAGAAGTTGGCTAGAAG | Amplification of 1316 DHA with an overlapping sequence for *mNeonGreen* |
| 1316 DHA Rev | GCGCATAGTGCTAGTCTTTTCTCC | |
| 1406 UHA For | GTTGGTATTCTCGATAGGCAGC | Amplification of 1406 UHA with an overlapping sequence for *TDH3p* |
| 1406 UHA Rev | TATTCTTTGAAATGGCAGTATTGATAATGAC<br>CCATCAGAACCGTAAACCTTG | |
| 1406 *TDH3p* For | GAAAGCGCCAAGGTTTACGGTTCTGATGG<br>GTCATTATCAATACTGCCATTTCAAAG | Amplification of promoter *TDH3p* with overlapping sequences for 1406 UHA and *mNeonGreen* |
| 1406 *mNeonGreen* Rev | AAGGATACTTCAAGACTAGATTCCCCCCTG<br>CGTTCAGGGTAATATATTTTAACCGCCG | Amplification of *mNeonGreen* with overlapping sequences for *TDH3p* and 1406 DHA |

| | | |
|---|---|---|
| 1406 DHA For | GTCGGCGGTTAAAATATATTACCCTGAACG<br>CAGGGGGGAATCTAGTCTTG | Amplification of 1406 DHA with an overlapping sequence for *mNeonGreen* |
| 1406 DHA Rev | GCGTCCTTATCGAAAGGAAC | |
| 1531 UHA For | GACTGCCTCTTGATGTTATGCCA | Amplification of 1531 UHA with an overlapping sequence for *TDH3p* |
| 1531 UHA Rev | TATTCTTTGAAATGGCAGTATTGATAATGAG<br>AAAGTTGCCGAGGCCAAATG | |
| 1531 *TDH3p* For | TATTTTCTCCATTTGGCCTCGGCAACTTTCT<br>CATTATCAATACTGCCATTTCAAAG | Amplification of promoter *TDH3p* with overlapping sequences for 1531 UHA and *mNeonGreen* |
| 1531 *mNeonGreen* Rev | ACGTAGATCGGTATATACGTTCAAGCCCCC<br>CGTTCAGGGTAATATATTTTAACCGCCG | Amplification of *mNeonGreen* with overlapping sequences for *TDH3p* and 1531 DHA |
| 1531 DHA For | GTCGGCGGTTAAAATATATTACCCTGAACG<br>GGGGGCTTGAACGTATATACC | Amplification of 1531 DHA with an overlapping sequence for *mNeonGreen* |
| 1531 DHA Rev | GGATGGCAGAACCGATACTAATG | |
| 1603 UHA For | GGCTATGGTGGTGATGTCTG | Amplification of 1603 UHA with an overlapping sequence for *TDH3p* |
| 1603 UHA Rev | TATTCTTTGAAATGGCAGTATTGATAATGAG<br>AGGAAAAAAAAACAGTTGTACATTGG | |
| 1603 *TDH3p* For | GTTACCAATGTACAACTGTTTTTTTTTCCTCT<br>CATTATCAATACTGCCATTTCAAAG | Amplification of promoter *TDH3p* with overlapping sequences for 1603 UHA and *mNeonGreen* |
| 1603 *mNeonGreen* Rev | AAAAAGCTCGTGAATACAGCAAGAACGAA<br>GCGTTCAGGGTAATATATTTTAACCGCCG | Amplification of *mNeonGreen* with overlapping sequences for *TDH3p* and 1603 DHA |
| 1603 DHA For | GTCGGCGGTTAAAATATATTACCCTGAACG<br>CTTCGTTCTTGCTGTATTCACG | Amplification of 1603 DHA with an overlapping sequence for *mNeonGreen* |
| 1603 DHA Rev | CTGTCTCCGCTATGTCAGTTAC | |
| 1531 UHA For | GACTGCCTCTTGATGTTATGCCA | Amplification of 1531 UHA with an overlapping sequence for *TDH3p* |
| 1531 UHA Rev | TATTCTTTGAAATGGCAGTATTGATAATGAG<br>AAAGTTGCCGAGGCCAAATG | |
| 1531 *TDH3p* For | TATTTTCTCCATTTGGCCTCGGCAACTTTCT<br>CATTATCAATACTGCCATTTCAAAG | Amplification of promoter *TDH3p* with overlapping sequences for 1531 UHA and *mCherry* |
| *TDH3p* (*mCherry*) Rev | CATGTTATCCTCCTCGCCCTTGCTCACCAT<br>TTTGTTTGTTTATGTGTGTTTATTCGAAAC | |
| 1531 *mCherry* For | GTTTCGAATAAACACACATAAACAAACAAA<br>ATGGTGAGCAAGGGCGAGG | Amplification of *mCherry* with overlapping sequences for *TDH3p* and 1603 DHA |
| 1531 *mCherry* Rev | ACGTAGATCGGTATATACGTTCAAGCCCCC<br>GGCTGGGAAGCATATTTGAGAAG | |
| 1531 DHA For | GCCGCATCTTCTCAAATATGCTTCCCAGCC<br>GGGGGCTTGAACGTATATACC | Amplification of 1531 DHA with an overlapping sequence for *mCherry* |
| 1531 DHA Rev | GGATGGCAGAACCGATACTAATG | |

* Black nucleotides represent the annealing part; red nucleotides represent the overlapping part of the primers.

** *TDH3p* Rev and *mNeonGreen* For primers were the same for all regions

**Table S4:** The primer list for multiple integrations of the β–carotene pathway genes and for colony PCR to control integrations

| **Name** | **Sequence 5' to 3'** | **Purpose** |
|---|---|---|
| 1406 UHA For | GTTGGTATTCTCGATAGGCAGC | Amplification of 1406 UHA with an overlapping sequence for *TDH3p* |
| 1406 UHA Rev | TATTCTTTGAAATGGCAGTATTGATAATGACCCATCAGAACCGTAAACCTTG | |
| 1406 *TDH3p* For | GAAAGCGCCAAGGTTTACGGTTCTGATGGGTCATTATCAATACTGCCATTTCAAAG | Amplification of promoter *TDH3p* with overlapping sequences for 1406 UHA and *CrtE* |
| *TDH3p* (*CrtE*) Rev | GATAGCGGTCAAGATGTTAGCGTAGTCCATTTTGTTTGTTTATGTGTGTTTATTCGAAAC | |
| *CrtE* For | GTTTCGAATAAACACACATAAACAAACAAAATGGACTACGCTAACATCTTGACCG | Amplification of *CrtE* with overlapping sequences for *TDH3p and* 1406 DHA |
| *CrtE* Rev | AAGGATACTTCAAGACTAGATTCCCCCCTGCGTTCAGGGTAATATATTTTAACCGCCG | |
| 1406 DHA For | GTCGGCGGTTAAAATATATTACCCTGAACGCAGGGGGGAATCTAGTCTTG | Amplification of 1406 DHA with an overlapping sequence for *CrtE* |
| 1406 DHA Rev | GCGTCCTTATCGAAAGGAAC | |
| 1531 UHA For | GACTGCCTCTTGATGTTATGCCA | Amplification of 1531 UHA with an overlapping sequence for *TDH3p* |
| 1531 UHA Rev | TATTCTTTGAAATGGCAGTATTGATAATGAGAAAGTTGCCGAGGCCAAATG | |
| 1531 *TDH3p* For | TATTTTCTCCATTTGGCCTCGGCAACTTTCTCATTATCAATACTGCCATTTCAAAG | Amplification of promoter *TDH3p* with overlapping sequences for 1531 UHA and *CrtYB* |
| *TDH3p* (*CrtYB*) Rev | GTGGATTTGGTAGTAAGCCAAAGCGGTCATTTTGTTTGTTTATGTGTGTTTATTCGAAAC | |
| *CrtYB* For | GTTTCGAATAAACACACATAAACAAACAAAATGACCGCTTTGGCTTACTAC | Amplification of *CrtYB* with overlapping sequences for *TDH3p and* 1531 DHA |
| *CrtYB* Rev | ACGTAGATCGGTATATACGTTCAAGCCCCCCGTTCAGGGTAATATATTTTAACCGCCG | |
| 1531 DHA For | GTCGGCGGTTAAAATATATTACCCTGAACGGGGGGCTTGAACGTATATACC | Amplification of 1531 DHA with an overlapping sequence for *CrtYB* |
| 1531 DHA Rev | GGATGGCAGAACCGATACTAATG | |
| 1603 UHA For | GGCTATGGTGGTGATGTCTG | Amplification of 1603 UHA with an overlapping sequence for *TDH3p* |
| 1603 UHA Rev | TATTCTTTGAAATGGCAGTATTGATAATGAGAGGAAAAAAAACAGTTGTACATTGG | |
| 1603 *TDH3p* For | GTTACCAATGTACAACTGTTTTTTTTCCTCTCATTATCAATACTGCCATTTCAAAG | Amplification of promoter *TDH3p* with overlapping sequences for 1603 UHA and *CrtI* |
| *TDH3p* (*CrtI*) Rev | TGGCTTGTCTTGGTCTTGTTCCTTACCCATTTTGTTTGTTTATGTGTGTTTATTCGAAAC | |
| *CrtI* For | GTTTCGAATAAACACACATAAACAAACAAAATGGGTAAGGAACAAGACCAAG | Amplification of *CrtI* with overlapping sequences for *TDH3p and* 1603 DHA |
| *CrtI* Rev | AAAAAGCTCGTGAATACAGCAAGAACGAAGCGTTCAGGGTAATATATTTTAACCGCCG | |
| 1603 DHA For | GTCGGCGGTTAAAATATATTACCCTGAACGCTTCGTTCTTGCTGTATTCACG | Amplification of 1603 DHA with an overlapping sequence for *CrtI* |
| 1603 DHA Rev | CTGTCTCCGCTATGTCAGTTAC | |
| *CrtE* CDS Rev | CCTCATCAAGATTGCTTTATGCCACTCGAGctaCAATGGGATGTCAGCCAACTTCTTC | OE PCR primers for assembly of *CrtE* CDS and terminator *TDH1t* |
| *TDH1t* (*CrtE*) For | TTGAAGAAGTTGGCTGACATCCCATTGtagCTCGAGTGGCATAAAGCAATCTTG | |
| *CrtYB* CDS Rev | CCTCATCAAGATTGCTTTATGCCACTCGAGctaTTGACCTTCCCAACCAGACA | OE PCR primers for assembly of *CrtYB* CDS and terminator *TDH1t* |
| *TDH1t* (*CrtYB*) For | GTTGTTATGTCTGGTTGGGAAGGTCAAtagCTCGAGTGGCATAAAGCAATCTTG | |

| | | |
|---|---|---|
| *CrtI* CDS Rev | CCTCATCAAGATTGCTTTATGCCACTCGAGctaG AAAGCCAAAACACCAACAGATC | OE PCR primers for assembly of *CrtI* CDS and terminator *TDH1t* |
| *TDH1t* (*CrtI*) For | GCTCGATCTGTTGGTGTTTTGGCTTTCtagCTCG AGTGGCATAAAGCAATCTTG | |
| *CrtE* Col PCR For | GAAGAAGTTGGCTGACATCC | Colony PCR for *CrtE* integration |
| 1406 Col PCR Rev | GCGTCCTTATCGAAAGGAAC | |
| *CrtYB* Col PCR For | GTTGTTATGTCTGGTTGGGAAGG | Colony PCR for *CrtE* integration |
| 1531 Col PCR Rev | GGATGGCAGAACCGATACTAATG | |
| *CrtI* Col PCR For | GCTCGATCTGTTGGTGTTTTG | Colony PCR for *CrtE* integration |
| 1603 Col PCR Rev | CTGTCTCCGCTATGTCAGTTAC | |

* Black nucleotides represent the annealing part; red nucleotides represent the overlapping part of the primers.
** The gene names in the parentheses show that the corresponding primer contains overlapping parts for the gene in the parenthesis.

**Table S5:** The primer and crRNA list for genomic deletions

| Name | Sequence 5' to 3' | Purpose |
|---|---|---|
| crRNA GAL80 | GATGAGCGTGGTAACCGATT ***GGG*** | 20 bp crRNA for *GAL80* deletion |
| GAL80 DEL UHA For | GCCTGTCTACAGGATAAAGACG | Amplification of UHA for *GAL80* deletion with an overlapping sequence for DHA |
| GAL80 DEL UHA Rev | ACTGGGGGCCAAGCACAGGGCAAGATGCTTGA CGGGAGTGGAAAGAACGG | |
| GAL80 DEL DHA For | AGTTGGTTTCCCGTTCTTTCCACTCCCGTCAAG CATCTTGCCCTGTGCTTG | Amplification of DHA for *GAL80* deletion with an overlapping sequence for UHA |
| GAL80 DEL DHA Rev | CCAGCAAAAATATGACCCCC | |
| crRNA DIT1 | CCTTGTCAAGATATATCCAC ***CGG*** | 20 bp crRNA for *DIT1* deletion |
| DIT1 DEL UHA For | GAGTCCCTGGAAGGAAAATTATTG | Amplification of UHA for *DIT1* deletion with an overlapping sequence for DHA |
| DIT1 DEL UHA Rev | TATTCCCCCTCTGTTAAATGGAATTGTGTGGCG GAGGAGCACAATTTATG | |
| DIT1 DEL DHA For | TAAATTTTCACATAAATTGTGCTCCTCCGCCACA CAATTCCATTTAACAGAGG | Amplification of DHA for *DIT1* deletion with an overlapping sequence for UHA |
| DIT1 DEL DHA Rev | CTGATGCCTCAAGATTTAACC | |

* Black nucleotides represent the annealing part; red nucleotides represent the overlapping part of the primers.
** The **bold** and *italic* sequences show the PAM sequences of corresponding crRNAs

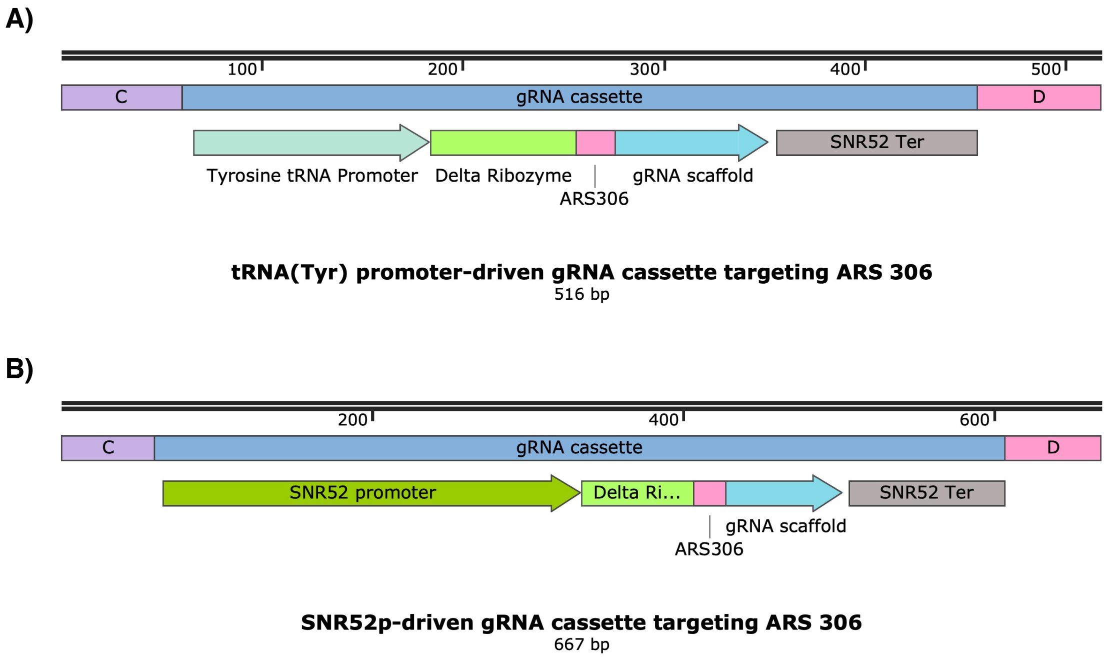


**Figure S1:** The sequence maps of gRNA cassettes driven by different promoters. **A)** The gRNA cassette is expressed through the $tRNA^{Tyr}$ promoter. It targets ARS 306 on chromosome III. The total length of is 516 bp. **B)** The gRNA cassette is expressed through *SNR52p*. It targets ARS 306 on chromosome III. The total size is 667 bp. C and D represent the 60 bp ortholog fragments used for *in vivo* DNA assembly with the other plasmid parts. The maps were illustrated using SnapGene®. [2]

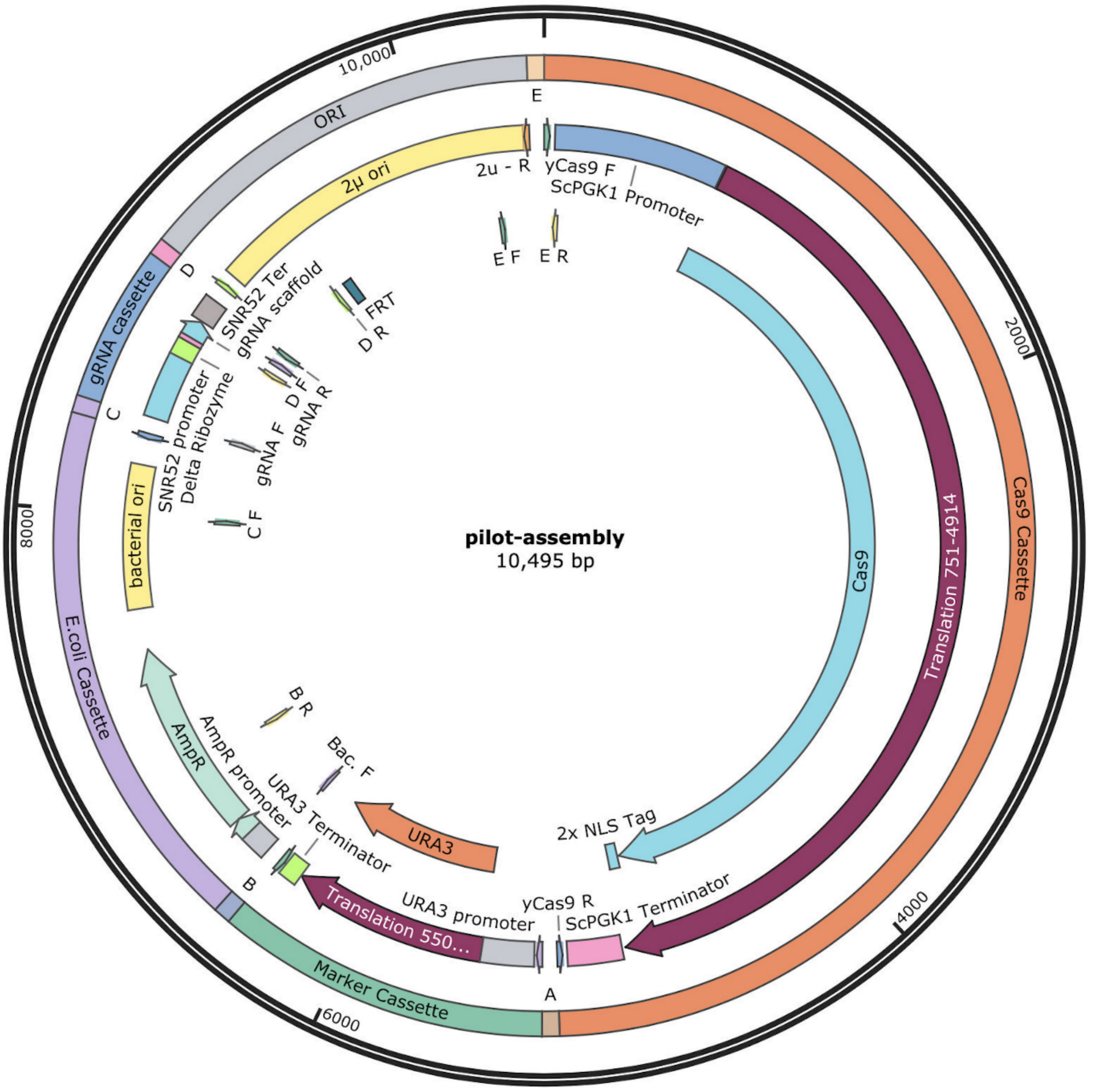


**Figure S2:** The plasmid map of correct *in vivo* DNA assembly. The short arrows containing F (forward) or R (reverse) suffixes represent primers, while A, B, C, D, and E represent 60 bp synthetic fragments. The primers contain 60 bp synthetic overlapping regions with the adjacent fragment. The system was designed using Benchling.[3] The map was illustrated using SnapGene®. [2]

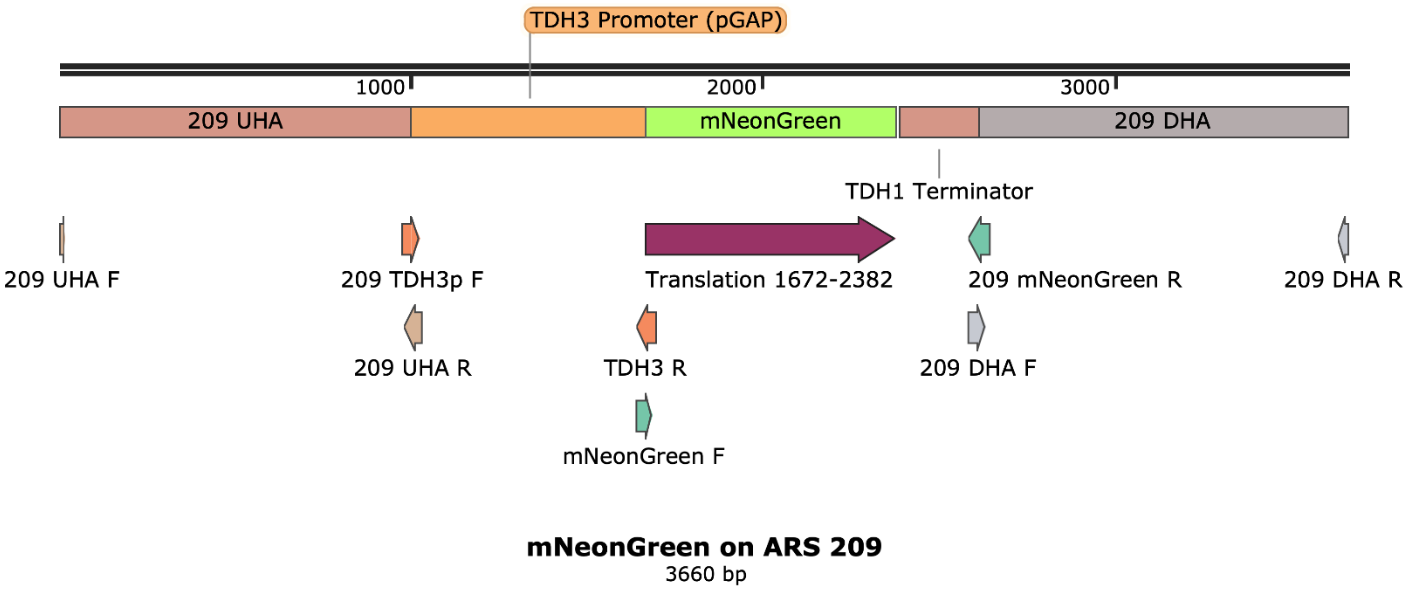


**Figure S3:** Genomic integration of *mNeonGreen* after *in vivo* DNA assembly. ARS 209 region is shown in the figure as an example. A similar design was also used for the other regions. Arrows represent primers. Longer primers contain overlapping regions with the adjacent fragment. The system was designed using Benchling.[3] The map was illustrated using SnapGene®. [2] F: forward primer, R: reverse primer

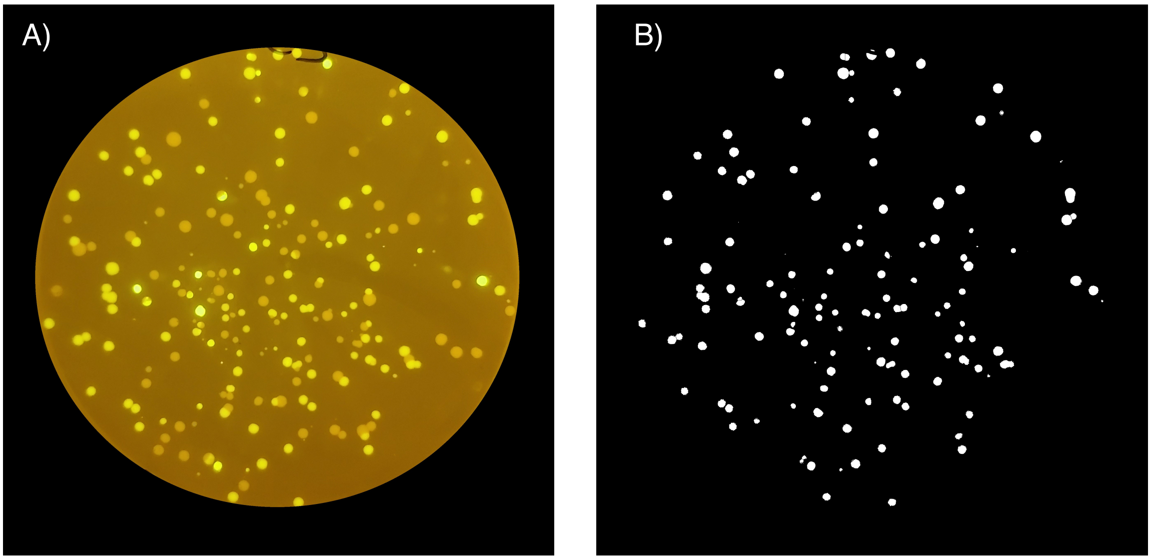


**Figure S4:** Automated counting of mNeonGreen expressing positive colonies using ImageJ and its Colony Counter plug-in. [4,5] **A)** The first image taken on a blue-LED transilluminator shows both fluorescent mNeonGreen-expressing and non-fluorescent false-positive colonies. **B)** The processed image to automatically select the positive colonies. The image was converted to a 16-bit format, the background and white negative colonies were removed using color threshold, the merged colonies were segmented to obtain individual colonies.

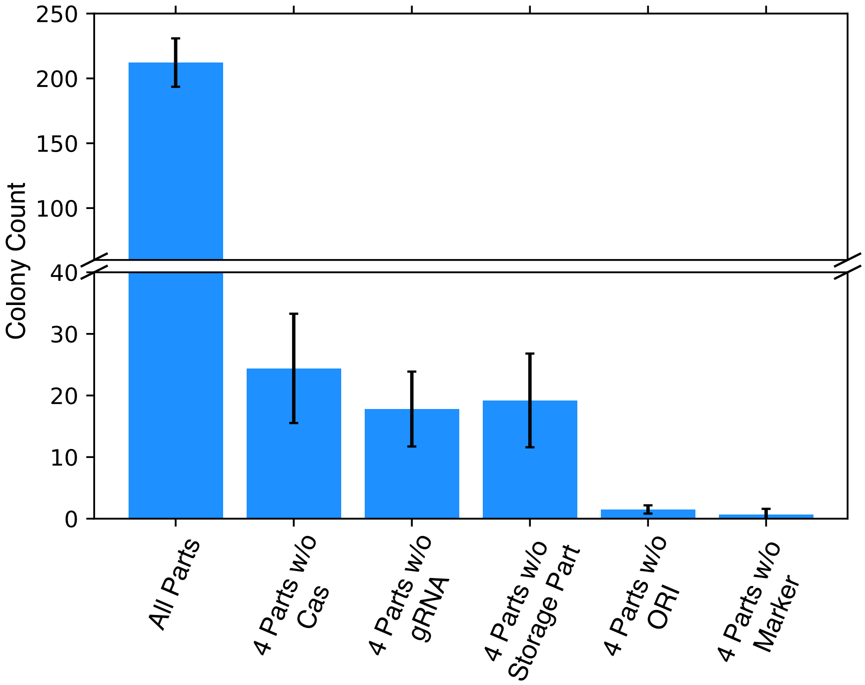


**Figure S5:** Colony numbers following transformation with different plasmid part combinations. The same transformation conditions were used for each combination. More than 200 colonies were obtained when all parts were used as they had overlapping sequences for *in vivo* assembly via a homology-directed repair mechanism. The error bars represent the standard deviations of three independent replicates.

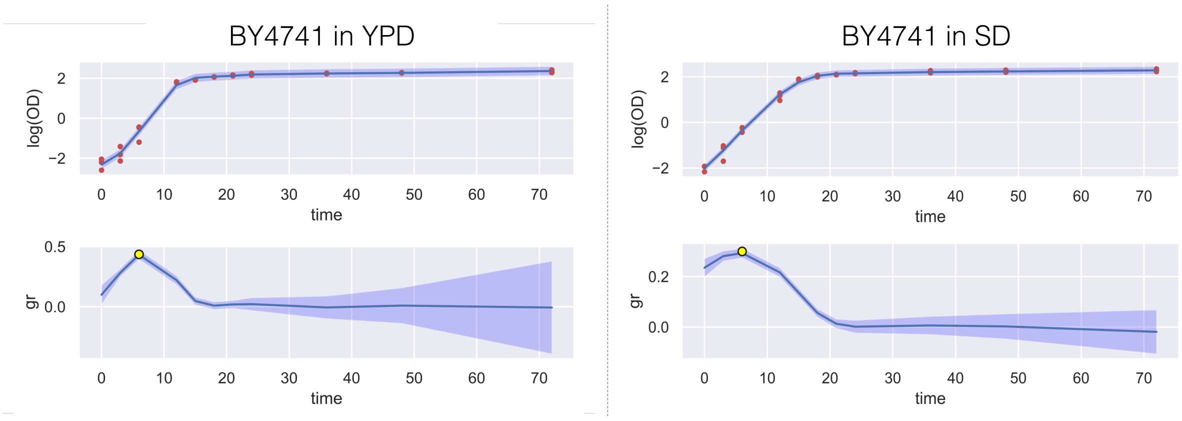


**Figure S6:** Logarithmic $OD_{600}$ (log(OD)) and the growth rate (gr) of the parental strain, BY4741, in YPD and SD media, respectively, over 72 hours (time). The yellow dots represent the maximum point of the curves. The standard deviations of three independent colonies are shown by shading.

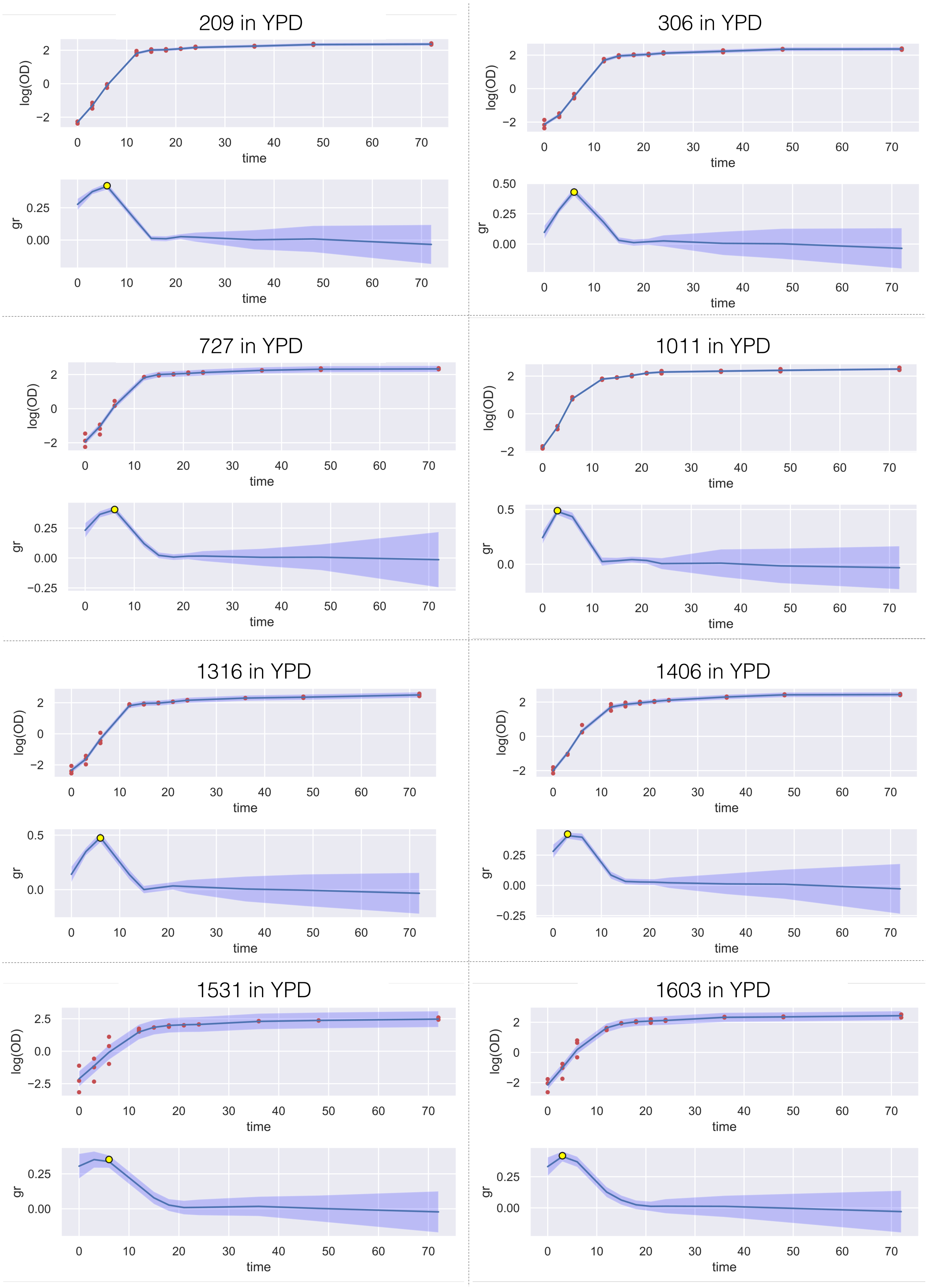


**Figure S7:** Logarithmic $OD_{600}$ (log(OD)) and growth rates (gr) of *mNeonGreen* integrated strains in YPD media over 72 hours (time). The yellow dots represent the maximum point of the curves. The standard deviations of three independent colonies are shown by shading.

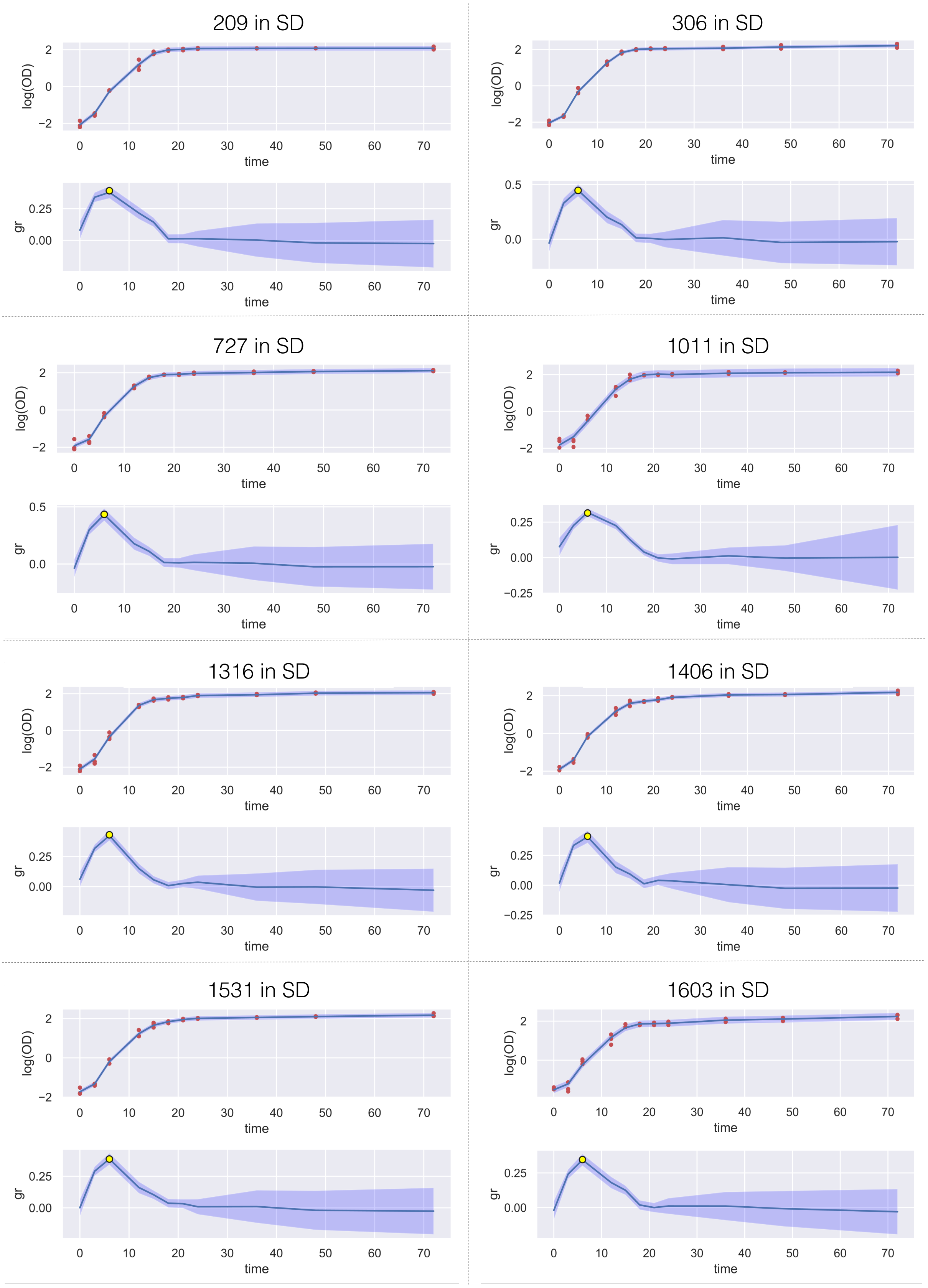


**Figure S8:** Logarithmic $OD_{600}$ (log(OD)) and growth rates (gr) of *mNeonGreen* integrated strains in SD media over 72 hours (time). The yellow dots represent the maximum point of the curves. The standard deviations of three independent colonies are shown by shading.

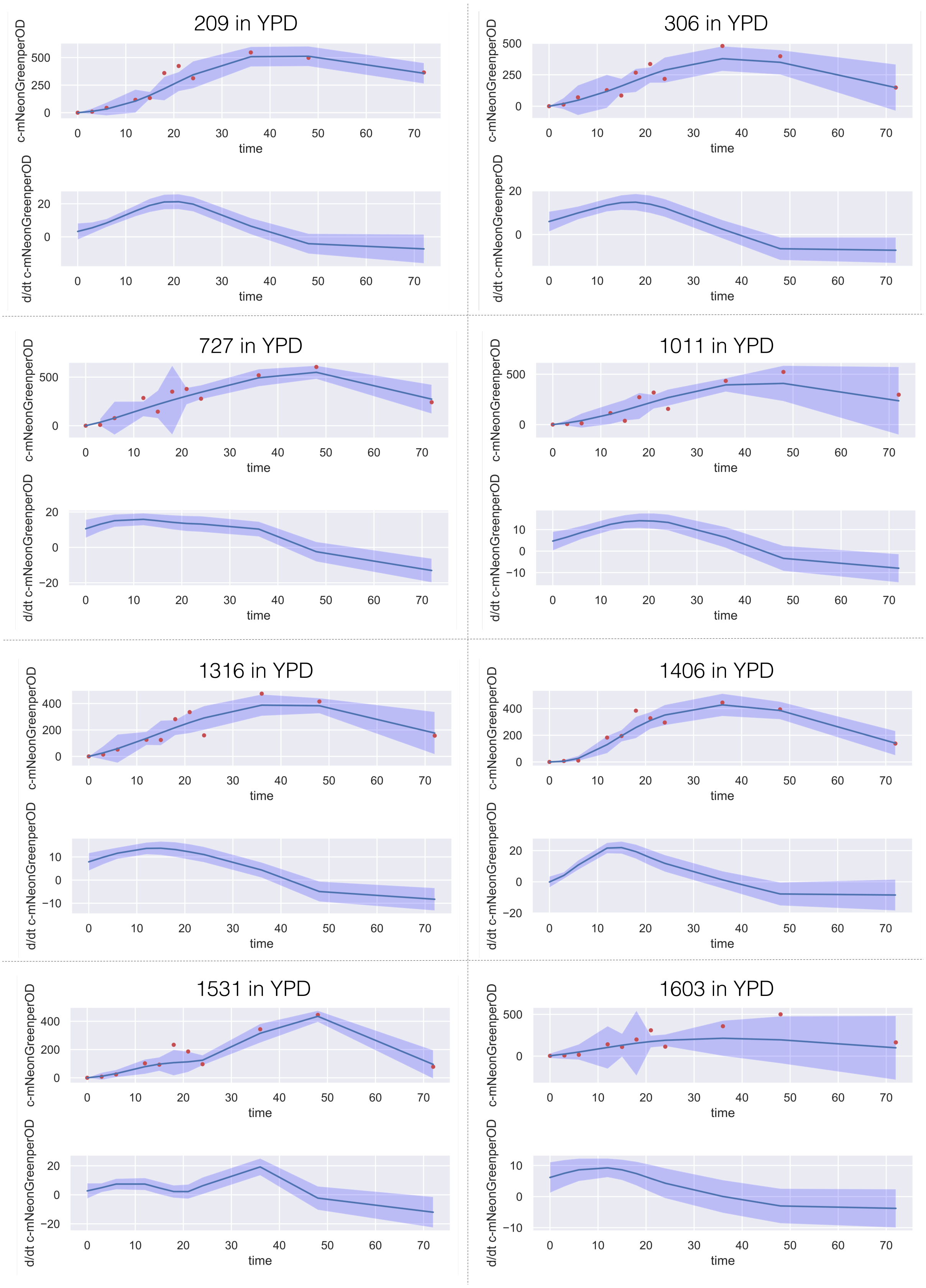


**Figure S9:** Corrected mNeonGreen expression per OD (c-mNeonGreenperOD) and time-derivative estimations of mNeonGreen expressions (d/dt -mNeonGreenperOD) of *mNeonGreen* integrated strains over 72 hours (time) in YPD media. The expressions are shown in the relative fluorescence unit (RFU). The standard deviations of three independent colonies are shown by shading.

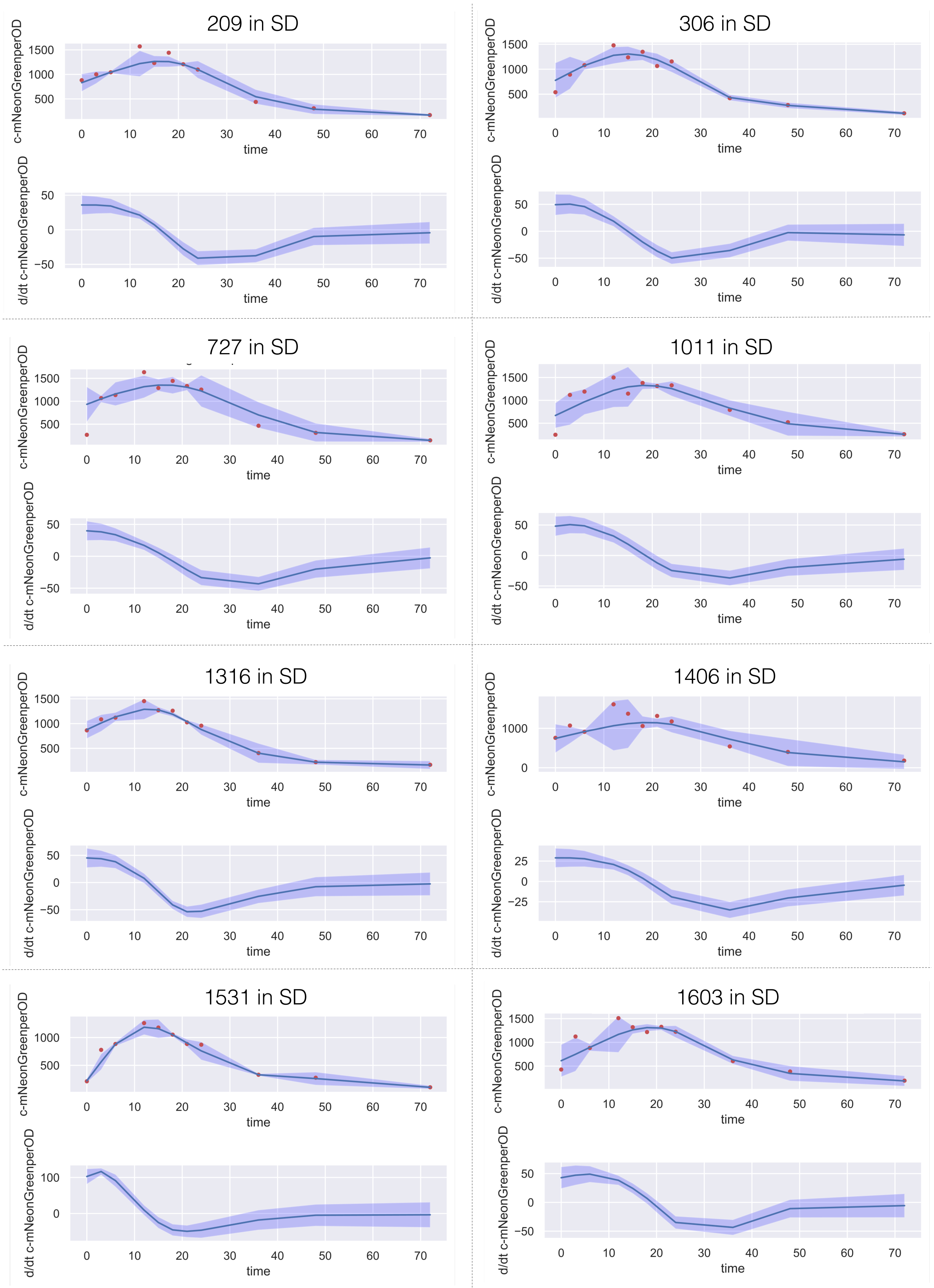


**Figure S10:** Corrected mNeonGreen expression per OD (c-mNeonGreenperOD) and time-derivative estimations of mNeonGreen expressions (d/dt -mNeonGreenperOD) of *mNeonGreen* integrated strains over 72 hours (time) in SD media. The expressions are shown in the relative fluorescence unit (RFU). The standard deviations of three independent colonies are shown by shading.

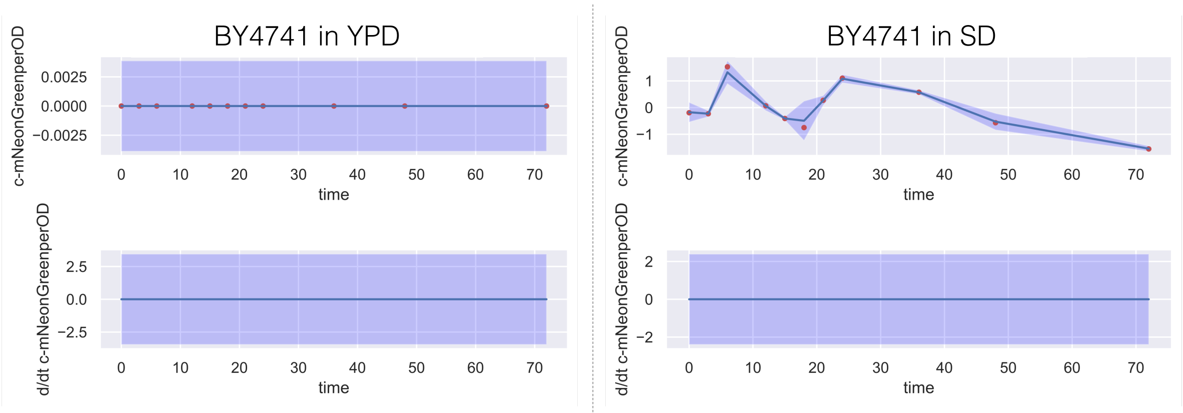


**Figure S11:** Corrected mNeonGreen expression per OD (c-mNeonGreenperOD) and time-derivative estimations of mNeonGreen expressions (d/dt -mNeonGreenperOD) of parental strain, BY4741, over 72 hours (time). The expressions are shown in the relative fluorescence unit (RFU). The standard deviations of three independent colonies are shown by shading.

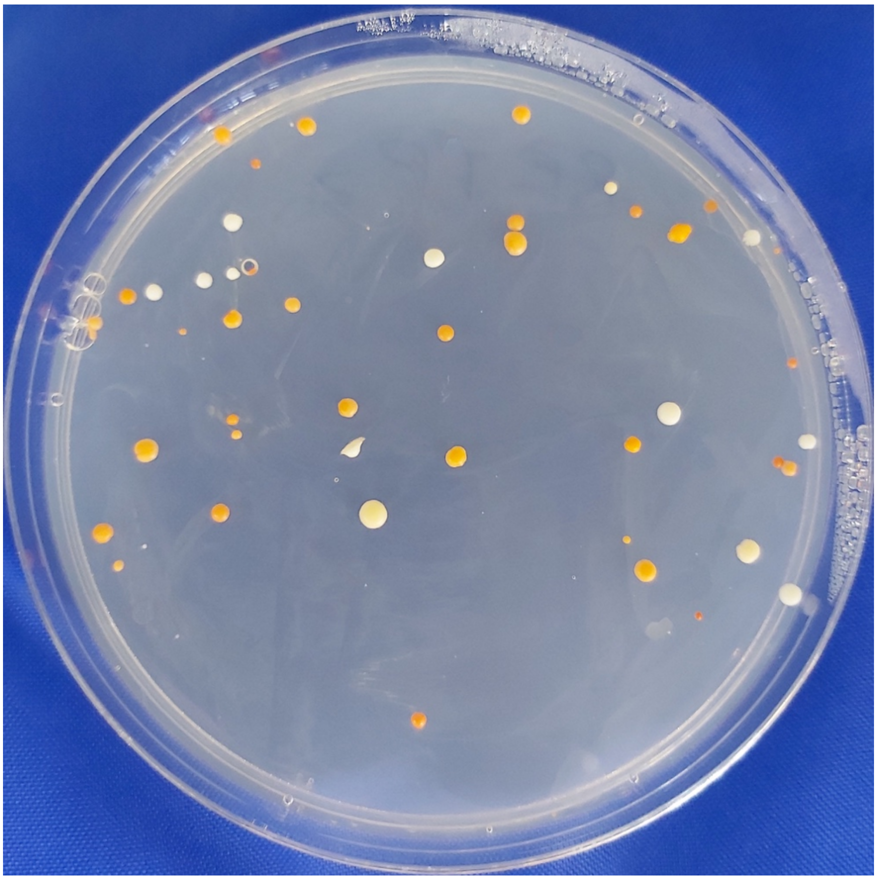

**Figure S12:** Multi-gene integration of three heterologous genes from the β-carotene pathway onto the ARS1531 region using ACtivE. The orange colonies show the β-carotene production by the correctly integrated genes, whereas the white or yellowish colonies have the missing gene(s) in their genome.
